# Navigating the structural landscape of *de novo* α–helical bundles

**DOI:** 10.1101/503698

**Authors:** Guto G. Rhys, Christopher W. Wood, Joseph L. Beesley, Nathan R. Zaccai, Antony J. Burton, R. Leo Brady, Andrew R. Thomson, Derek N. Woolfson

## Abstract

The association of amphipathic *α* helices in water leads to α-helical-bundle protein structures. However, the driving force for this—the hydrophobic effect—is not specific and does not define the number or the orientation of helices in the associated state. Rather, this is achieved through deeper sequence-to-structure relationships, which are increasingly being discerned. For example, for one structurally extreme but nevertheless ubiquitous class of bundle—the α-helical coiled coils—relationships have been established that discriminate between all-parallel dimers, trimers and tetramers. Association states above this are known, as are antiparallel and mixed arrangements of the helices. However, these alternative states are less-well understood. Here, we describe a synthetic-peptide system that switches between parallel hexamers and various up-down-up-down tetramers in response to single-amino-acid changes and solution conditions. The main accessible states of each peptide variant are characterized fully in solution and, in most cases, to high-resolution X-ray crystal structures. Analysis and inspection of these structures helps rationalize the different states formed. This navigation of the structural landscape of α-helical coiled coils above the dimers and trimers that dominate in nature has allowed us to design rationally a well-defined and hyperstable antiparallel coiled-coil tetramer (apCC-Tet). This robust *de novo* protein provides another scaffold for further structural and functional designs in protein engineering and synthetic biology.

## INTRODUCTION

In nature, bundles of four α helices are common in protein structures and assemblies. These four-helix bundles perform a wide variety of functions including: acting as protein hormones and cytokines;^1, 2^ providing scaffolds for metal- and co-factor-binding to facilitate storage, redox and enzymatic functions;^3–5^ cementing protein-protein interactions that direct DNA binding, membrane fusion and other processes;^6–8^ serving as building blocks for viral capsids;^9^ and spanning membranes to perform signal transduction^10^ and transport functions.^11^

As such, four-helix bundles have become key targets and scaffolds for *de novo* protein design.^12–18^ Usually, these comprise amphipathic α helices. These helices assemble *via* their hydrophobic faces to form bundles with the hydrophobic side chains buried in a consolidated core. The associations can be of individual helices to form tetramers, of helical hairpins to give dimers, or intramolecularly within the same protein chain.^19^ However, and particularly for the intermolecular cases, alternative helical arrangements (*e.g.*, all parallel, up-down antiparallel) and even other oligomeric states are possible. Therefore, the specification of helix-helix interactions that direct towards a specified structure and away from unwanted alternatives is critical for successful design; these two aspects are known as positive and negative design, respectively. Ultimately, what is required are clear sequence-to-structure relationships and/or robust computational methods to augment the amphipathic α-helical sequences and specify a target four-helix bundle. Here we explore the structural plasticity between all-parallel and up-down-up-down antiparallel four-helix coiled-coil structures and some of the other competing states.

Four-helix bundles are one of the earliest examples of a recognized protein-structure motif.^20^ Since then, structures of other four-helix bundles have enabled analyses of features that define the fold. Weber and Salemme expand on previous work, including Crick’s model for packing in α-helical coiled coils,^21^ to show that structural similarities between four-helix bundles are a consequence of basic physical properties of the component helices.^22^ Presnell and Cohen present further examples of four-helix bundles using computational screening and visual inspection of over 300 protein structures.^23^ This screening identifies 20 putative structures based on buried surface area, but when the 20 are examined visually, some are excluded as they “do not appear to be well packed”, with helix crossing angles diverging from extant four-helix bundles. These examples of true four-helix bundles provide the basis for categorizing the fold by helix-crossing angles, with six categories emerging: square, splinter, x, unicornate, bicornate and splayed.^24^ The square class contains the previously identified four-helix bundles, where all helices are aligned, while the other classes have packing more similar to previously identified α-helical globules.^25^ The main differentiating factor between these structures is the number and regularity of interactions between helices.

Indeed, there is spectrum of side-chain-mediated helix-to-helix packing in four-helix bundles from few non-specific interactions to repeated regular interactions (Figure 1A).^26^ Short helices (< 14 residues) tend to be more free to associate in aligned or orthogonal configurations, resulting in a wide range of bundles.^25^ Whereas, longer helices (>= 14 residues) favor aligned arrangements, packing optimally as extended bundles with regular and repeating interactions.^27^ Extended bundles can be categorized further based on the predominant mode of helix packing, from less-specific ridges-into-grooves (RIG) to intimate knobs-into-holes (KIH) interactions.^21,28,29^ These packing modes are characteristics of globular and coiled-coil domains, respectively.

**Figure 1.**
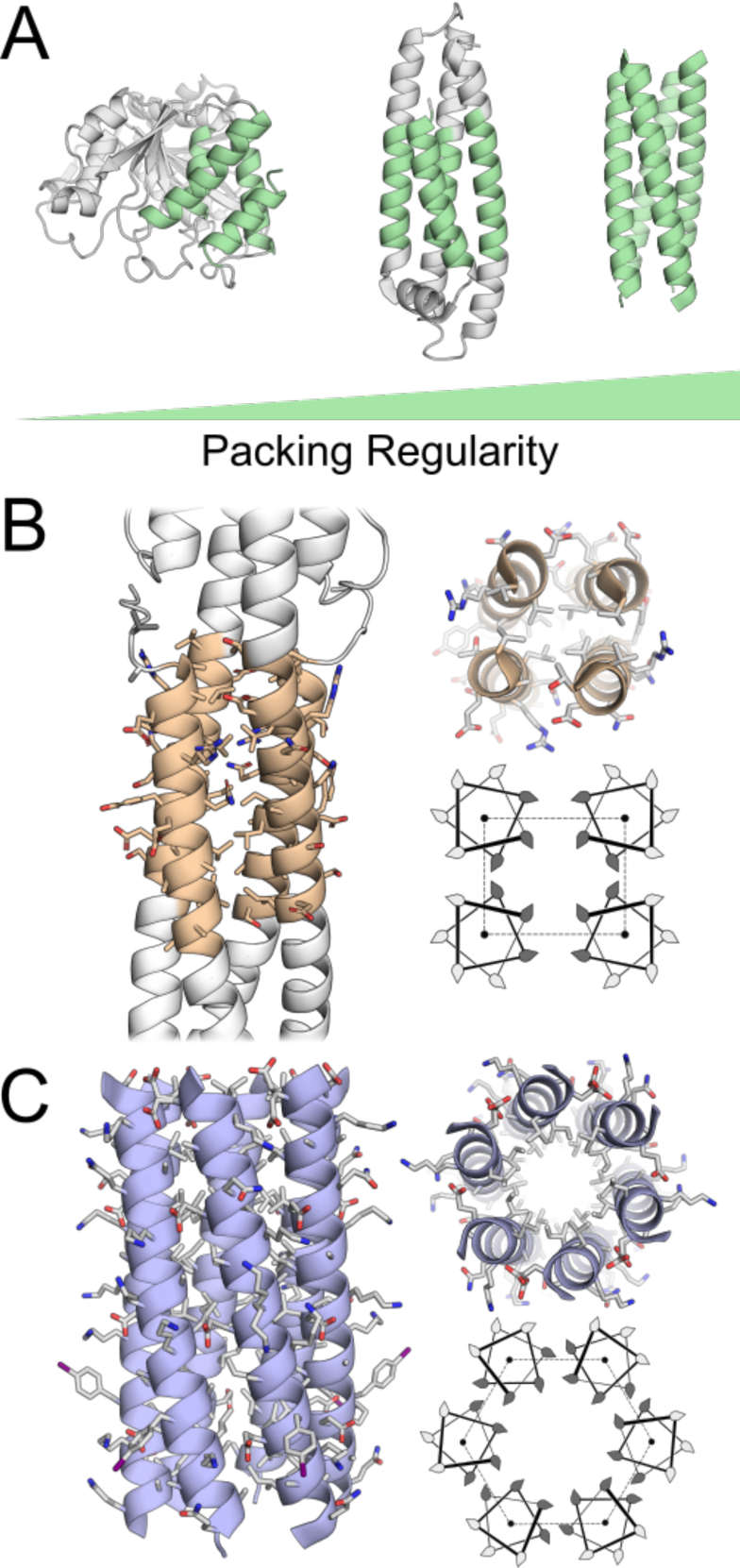
Four-helix bundles adopt a range of topologies with irregular to highly regular packing. From left to right, the structures are for a region of 3-isopropylmalate dehydrogenase (PDB ID code: 1IPD) in the x class with only pairs of helices aligned; apolipoprotein E3 (PDB ID code:1LPE) in the square class, where all are helices aligned and interact *via* ridges-into-grooves packing; and CCTet (PDB ID code: 3R4A) also in the square class but with knobs-into-holes helix-helix interactions. (B) The HAMP domain of TSR/AF1503, which forms an alacoil (left and top right, PDB ID code: 3ZX6), with the corresponding helical wheel showing helix packing (bottom right). (C) A designed hexameric coiled coil, CC-Hex (left and top right), also with a helical wheel of helix packing (bottom right).

Given their ubiquity and this level of understanding, four-helix bundles offer a fertile ground for the design and engineering of new proteins. DeGrado has pioneered this area:^30^ initially, designing small synthetic peptides of leucine, glutamate and lysine to create amphipathic helices,^31^ although these were later revealed to form larger globular bundles;^32^ then joining these together with proline-and-arginine-based loops to generate hairpins that dimerize;^33–36^ and expressing single-chain four-helix bundles that are monomeric and hyperstable.^19^ This bottom-up approach has led to sophisticated *de novo* four-helix bundles with ion-transportation and cofactor-binding properties.^37, 38^ Dutton and co-workers have also built four-helix bundle maquettes that incorporate heme to bind and transport oxygen.^39–41^ Similar scaffolds have been developed by others to introduce light-harvesting and enzyme-like functions.^42–44^

Hecht and co-workers have taken a different approach to designing four-helix bundles.^14^ This pioneers the use of binary patterns of hydrophobic (h) and polar (p) residues to define amphipathic helices linked into single chains by intervening turns.^45^ Rather than specifying the sequences any further, however, redundant codons are used to place multiple different residues at the h and p sites; *i.e.*, sequence libraries are generated to be compatible with the overall four-helix-bundle fold. Moreover, rather than actively selecting from or evolving these libraries, proteins that survive or operate in cells are picked out passively. In this way, the group have achieved stably folded and structured *de novo* four-helix bundles.^46, 47^ In turn, these have been endowed with functions such as heme binding,^48^ abilities to substitute for deleted endogenous proteins,^49, 50^ small-molecule binding,^51^ and reducing copper toxicity.^52^

With some exceptions,^53^ these *de novo* four-helix bundles tend to have sequences that promote RIG packing of helices. Thus, while structural data is limited, it is likely that most *de novo* four-helix bundles have irregular side-chain packing, falling in the middle of the spectrum of Figure 1A. For more-intimate KIH packing and coiled-coil formation, two general features are required: (i) heptad repeats of h and p residues, hpphppp (usually denoted *abcdefg*), or related repeats with 3, 4 spacings of h residues;^54^ and (ii) specific combinations of predominantly aliphatic hydrophobic residues at the *a* and *d* sites.^55^ Designed four-helix bundles that are coiled coils lie far to the right of the spectrum of Figure 1A. Early designs in this area elucidated the rules for the formation of parallel four-helix bundles and also for parallel dimers and trimers.^56, 57^ These rules have been embellished and used in rational *de novo* design of many parallel coiled coils.^55, 58^

Previously, we have described the rational design and complete characterization through to X-ray protein crystal structures of a basis set of all-parallel dimeric, trimeric and tetrameric coiled coils.^59^ Serendipitously, a simple permutation to the sequence repeat of the tetramer, CC-Tet, produces an entirely new coiled-coil assembly, namely an all-parallel hexamer, CC-Hex.^60^ This has a central accessible channel.^61^ Therefore, it is an α-helical barrel. The formation of the structure can be rationalized as the mutations to CC-Tet expanded the hydrophobic surfaces of the component helices allowing more of them to associate. Moreover, the introduction of complementary charged aspartic acid (Asp, D) and histidine (His, H) residues at the core position Leu-24 in the otherwise hydrophobic central channel of CC-Hex to render two peptides—CC-Hex-D24 and CC-Hex-H24—that complement to form a parallel A3B3-type heterohexamer. However, through a number of unpublished studies, we find that the CC-Hex scaffold is not completely robust and its oligomer state changes with other polar mutations. Subsequently, we have developed parametric computational design to deliver a series of α-helical barrels, including pentamers, new hexamers and a heptamer.^62^ These α-helical barrels are more-robust to mutation and serve as platforms for rational design to introduce new functions.^63, 64^

Herein, we return to the mutants of CC-Hex and explore structural plasticity in the coiled-coil structural landscape. This must be navigated to deliver robust *de novo* designs. Specifically, we find that whilst CC-Hex-L24E assembles as a stable parallel hexamer at low pH, it accesses a less-stable up-down-up-down antiparallel tetrameric coiled-coil state near neutral pH. Moreover, substituting positively charged lysine (Lys, K) or 2,4-diaminobutyric acid (Dab) residues at the L24 position gives the antiparallel tetramer. The X-ray crystal structures for this state guide the rational design of a robust and hyperstable antiparallel tetramer, apCC-Tet, which we characterize fully. This requires consideration of both the composition of the hydrophobic core and of interhelical salt-bridging in *en bloc* mutations of the original CC-Hex sequence. apCC-Tet provides an additional *de novo* protein fold that could be of use in protein design and engineering, materials science, and synthetic biology.

## RESULTS AND DISCUSSION

### Polar mutations at core sites of CC-Hex cause structural switches in solution

To probe how robust CC-Hex-based sequences were to forming α-helical barrels, we synthesized variants of CC-Hex with a mutation at position 24 and without the *C*-terminal AlaGly dipeptide. The 24th position is an *a* position of the heptad repeat and, therefore, contributes to the cores of dimeric – tetrameric coiled coils and points towards the lumens of α-helical barrels. The variants included: negatively charged side chains, L24D and L24E; positively charged side chains, L24H, L24Dab (L-2,4-diaminobutyric acid) and L24K; and a non-polar side chain, L24Nle (norleucine) (Table 1).

**Table 1.**
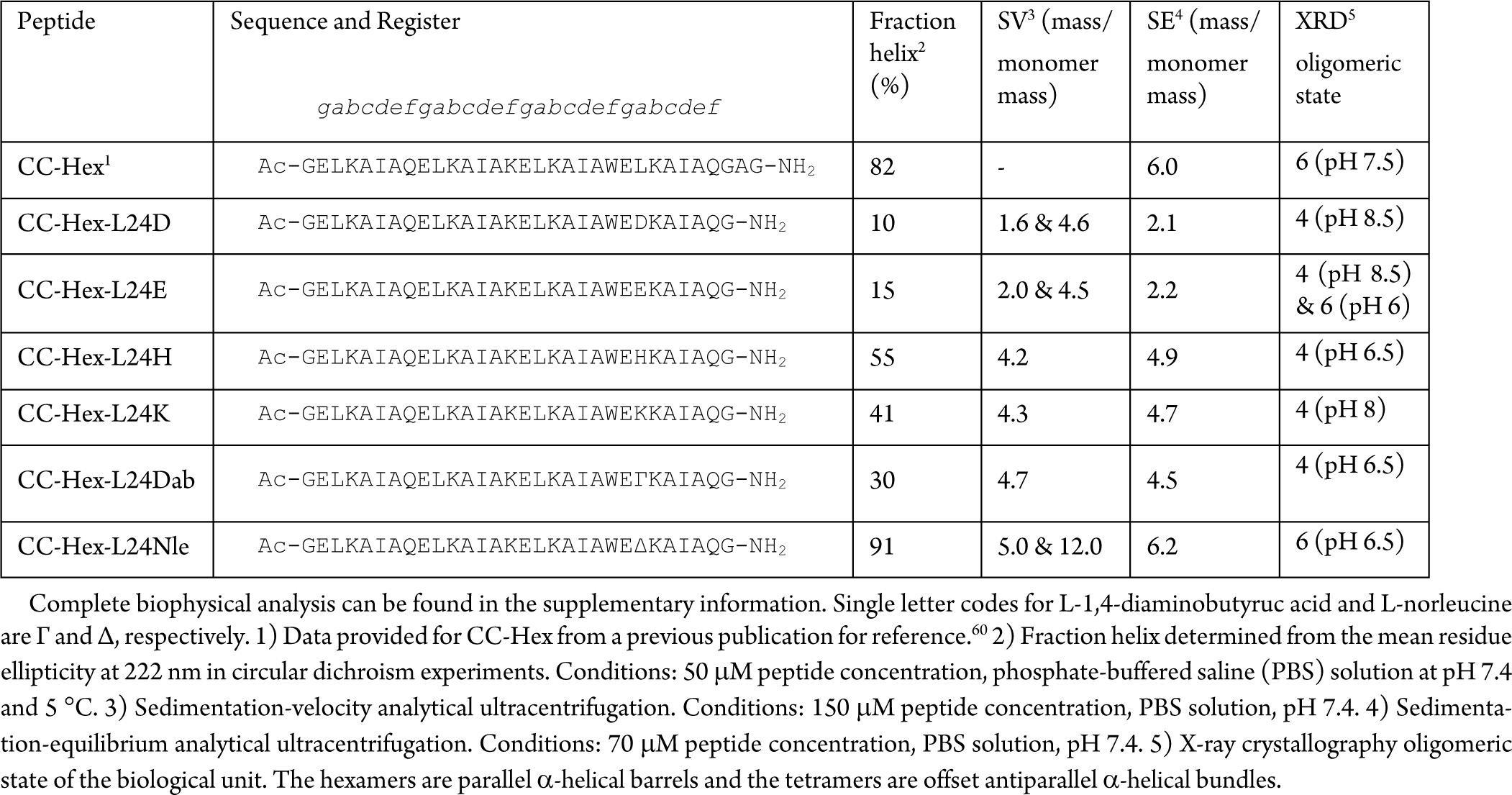
Sequences and summary of biophysical data for CC-Hex-L24 point mutants.

Circular dichroism (CD) spectroscopy of L24D and L24E at pH 7.4 showed that the peptides lost much of their helicity compared to the parent CC-Hex (Figure 2A). Analytical ultracentrifugation (AUC) experiments indicated that both variants formed a mixture of dimeric and tetrameric species rather than hexamers (Figure S6.1,S6.2; Table 1). For a given concentration, L24E was more α helical than L24D at neutral pH and, therefore, it was chosen for further biophysical characterization. Specifically, L24E was interrogated by CD spectroscopy and AUC over the pH range 3–7 (Figure 2B-D). At pH 3 and 4, the peptide was highly stable and did not fully unfold even by 90 °C. At pH 5 there was no significant loss of α helicity, but the peptide showed the start of a thermal unfolding transition. At pH 6 there was ≈30% drop in helicity and the peptide underwent full thermal denaturation. pH 7 saw a further drop in helicity and the peptide showed signs of cold denaturation. (N.B. This cold denaturation was typical of mutants confirmed as tetramers (Figures S4.1–4.4,S4.6)). Sedimentation-equilibrium AUC experiments revealed that from pH 3–7.4 L24E switched from hexameric to dimeric species (Figure 2D), whereas sedimentation velocity at pH 4 suggests multiple species (Figure S6.13). At pH 7.4, increasing the peptide concentration above 150 μM increased α helicity and shifted the equilibrium towards tetramers (Figures S5.1,S6.12).

**Figure 2.**
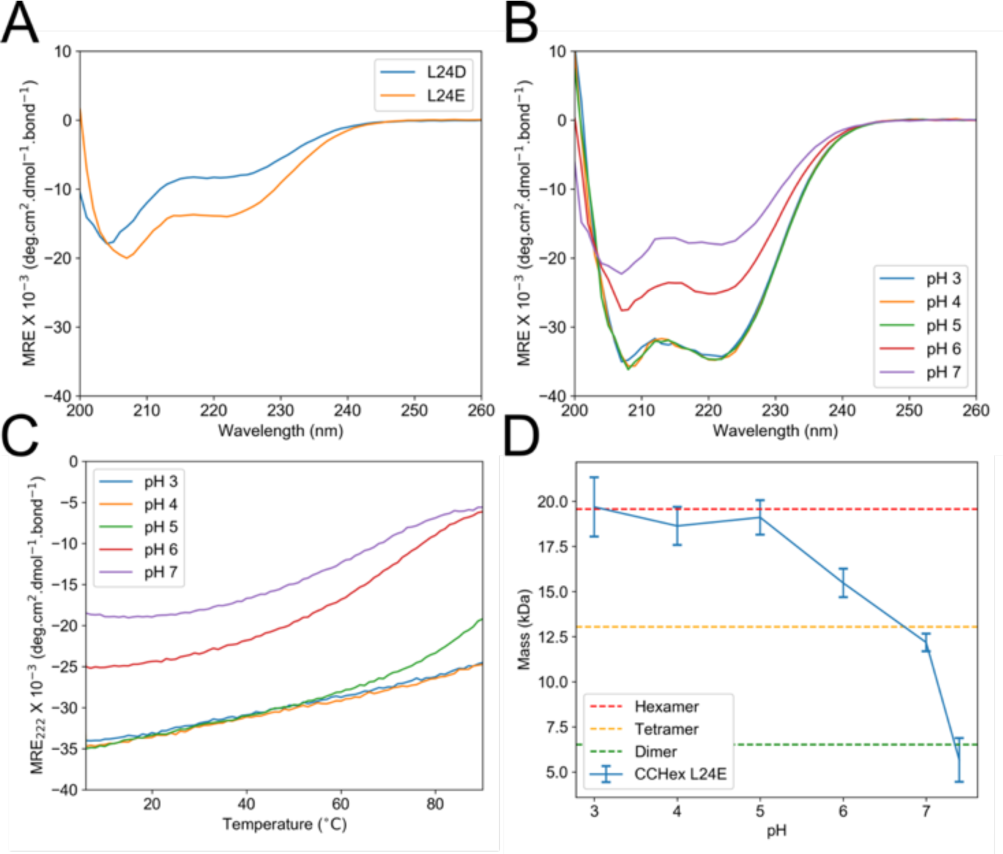
Solution-phase biophysical data for the CC-Hex-L24D and L24E variants. (A) Circular dichroism (CD) spectra for both peptides at 5 °C, 200 µM concentration, in phosphate-buffered saline (PBS) solution at pH 7.4. (B) CD spectra of CC-Hex-L24E at 5°C, 150 µM concentration in acidic to neutral conditions. (C) CD thermal denaturation profiles of CC-Hex-L24E from 5–90 °C in acidic to neutral conditions. (D) Molecular weight of CC-Hex-L24E in acidic to neutral conditions determined by sedimentation-equilibrium AUC experiments. Error bars show standard deviation (n=3).

To test if the tetramer resulted from charged residues at core sites generally, the positively charged variants, L24Dab, L24K and L24H (which has a pKa of 6.8) and the L24nLeu control were examined.^65^ Again, CD spectroscopy at pH 7.4 showed that L24H, L24K and L24Dab were significantly less α helical than the parent (Figures S4.3-4.4,S4.6). By SE AUC, L24K, L24Dab and L24H were not hexameric, with masses ∼5 x monomer mass (Figures S6.3-6.4,S6.6). By contrast, L24Nle was highly folded and hexameric in solution at pH 7.4; although an additional higher-order species was observed by sedimentation-velocity experiments accounting for 35% of the sample (Figures S4.5,S6.5).

Thus, the original CC-Hex is not robust to charged polar mutations in its hydrophobic channel, in contrast to the more robust computationally designed α-helical barrels.^63^ Moreover, at pH values where these residues are likely charged the structures are less helical, of lower thermal stability, and form lower oligomeric states.

### X-ray crystal structures reveal a broader accessible structural landscape

We obtained X-ray crystal structures for L24E, L24D, L24Dab, L24K, L24H and L24Nle. L24E crystallized in two forms that gave different structures: the previously documented hexameric blunt-ended barrel and an antiparallel tetramer (Figure 3A,C, respectively). The crystallization conditions for the two states were markedly different. The tetramer was only observed at pH 8.5, while the hexamer crystallized from several conditions from pH 5 – 6.5, as well from unbuffered solutions where the pH was low due to residual trifluoroacetic acid from peptide synthesis (pH < 2). L24D only crystallized as the antiparallel tetramer. However, previously we crystallized a hexameric form from unbuffered solution.^60^ L24K, L24Dab and L24H only crystallized as antiparallel tetramers, while L24Nle was only successfully crystallized in the hexameric form (Figure 3B); these structures were obtained in the pH range 6.5 – 8 (Table S2). For these four sequences, an inability to obtain other crystal forms does not necessarily mean that other states do not exist. Nevertheless, we note that the solution-phase biophysical data corroborate the observed crystal structures (Table 1).

**Figure 3.**
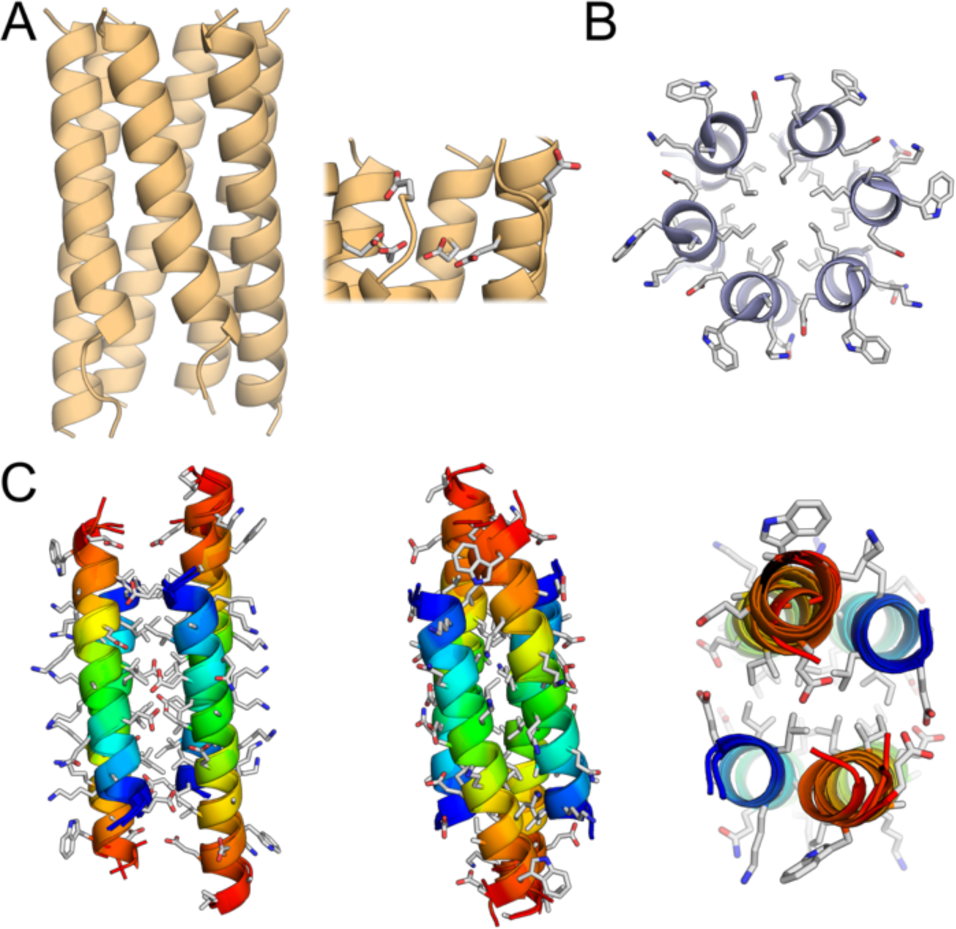
X-ray crystal structures of CC-Hex point mutants. (A) The hexameric crystal form of CC-Hex-L24E. (B) Slice through the structure of CC-Hex-L24Nle viewed from the *N* termini. (C) The overlaid backbones of the tetrameric forms of L24E, L24D, L24Dab, L24K and L24H with the side chains of L24E displayed for reference. The chains are colored in rainbow spectrum from the *N* termini (blue) to the *C* termini (red).

Comparing the low-pH hexameric structures of CC-Hex-D24 (a variant of CC-Hex-L24D with an AlaGly dipeptide extension at the *C* terminus, PDB ID code: 3R46) and L24E revealed the shorter aspartic acid could be accommodated within the core of a completely folded structure; whereas, glutamic acid could not be fully accommodated, as two of the six peptide chains frayed at the *C* terminus allowing the Glu side chains to extend into solvent (Figure 3A). Both L24E and L24K were more helical at neutral pH than the analogues with shorter side chains, L24D and L24Dab. It is possible that this is due to the higher helical propensity of the longer side chains, or more contributions from the methylene groups to helix-helix contacts.

The structures determined near neutral pH for L24D and L24E plus those for L24Dab, L24H and L24K are closely similar antiparallel four-helix coiled coils with backbone and all-atom RMSDs for residues 1-23 across of the whole set of 0.44 Å ± 0.12 Å and 1.13 Å ± 0.44 Å, respectively (Figure 3C). They have staggered rather than blunt-end arrangements of the helices, and the helices are frayed or disordered after the polar 24th residue. As a result, the cores are exclusively hydrophobic. The core has contributions from residues at *a* (Leu), *d* (Ile) and *e* (Ala) sites of the heptad repeats. This gives two distinct helix-helix interfaces: two “wide faces” centered on *d* = Ile (Figure 3C left) and two “narrow faces” centered on *e* = Ala (Figure 3C center). The Leu residues at *a* are directed towards central long axis of the bundle (Figure 3C right) reminiscent of complementary *x-da* layers observed in Alacoils (Figure 1B).^10^ In other words, the structures have oblate cross-sections. The two interfaces are flanked by pairs of the same amino acid from neighboring helices; *i.e., b:b* and *g:g*, respectively. As *b* = Lys and *g* = Glu, the narrow and wide faces present seams of positive and negative charge, respectively. Thus, the formation of the hydrophobic core overrides (i) the complete folding of each chain into α helices, and (ii) potential electrostatic repulsion between like residues at *b* and *g*.

Structural searches of PDBefold and CAME topsearch using the L24E tetramer identified the following: HAMP proteins (*e.g.* PDB ID code: 3ZX6; Cα RMSD = 1.2 Å); variants of the GCN4-p1 peptides (*e.g.* PDB ID code 1W5J; Cα RMSD = 1.3 Å); the ROP proteins (PDB ID code *e.g.* 1QX8; Cα RMSD = 1.6 Å); αTet (PDB ID code 6C52; Cα RMSD = 2.1 Å), a non-aggregating variant of a *de novo* designed cross-α amyloid-like structure; and di-Zn(II)-DF3l (PDB ID code 2KIK; Cα RMSD = 2.3 Å), an artificial protein designed for phenol oxidase activity. While all structures are antiparallel tetrameric bundles, only the HAMP proteins contain regions with oblate cross-section due to regular Alacoil. Nevertheless, the L24 variants are reminiscent of HexCoil-Ala (PDB ID code: 3S0R), a *de novo* designed protein fold that crystallizes as an antiparallel tetramer with Alacoil and wide interfaces.^66^ There are no polar, charged residues in the core of the HexCoil-Ala and the helices are fully folded and aligned blunt-ended. Furthermore, the core comprises *a, d* and *g* residues—as opposed to the L24 mutants having *a, d* and *e* residues—which results in the opposite-handedness topology.

### Blunt-ended and fully folded tetramers can be redesigned from the CC-Hex sequence

The structures of the polar mutants at L24 in the CC-Hex sequence demonstrate that another coiled-coil topology—*i.e.*, an up-down-up-down antiparallel tetramer—is accessible to our basis-set of *de novo* designs.^59,62,67^ However, the tetramers described thus far are not stable or fully folded. To explore how the fold might be optimized, we synthesized *en bloc* mutants of the CC-Hex sequence at the *a* & *d* and *g* & *b* sites (Table 2).

**Table 2.**
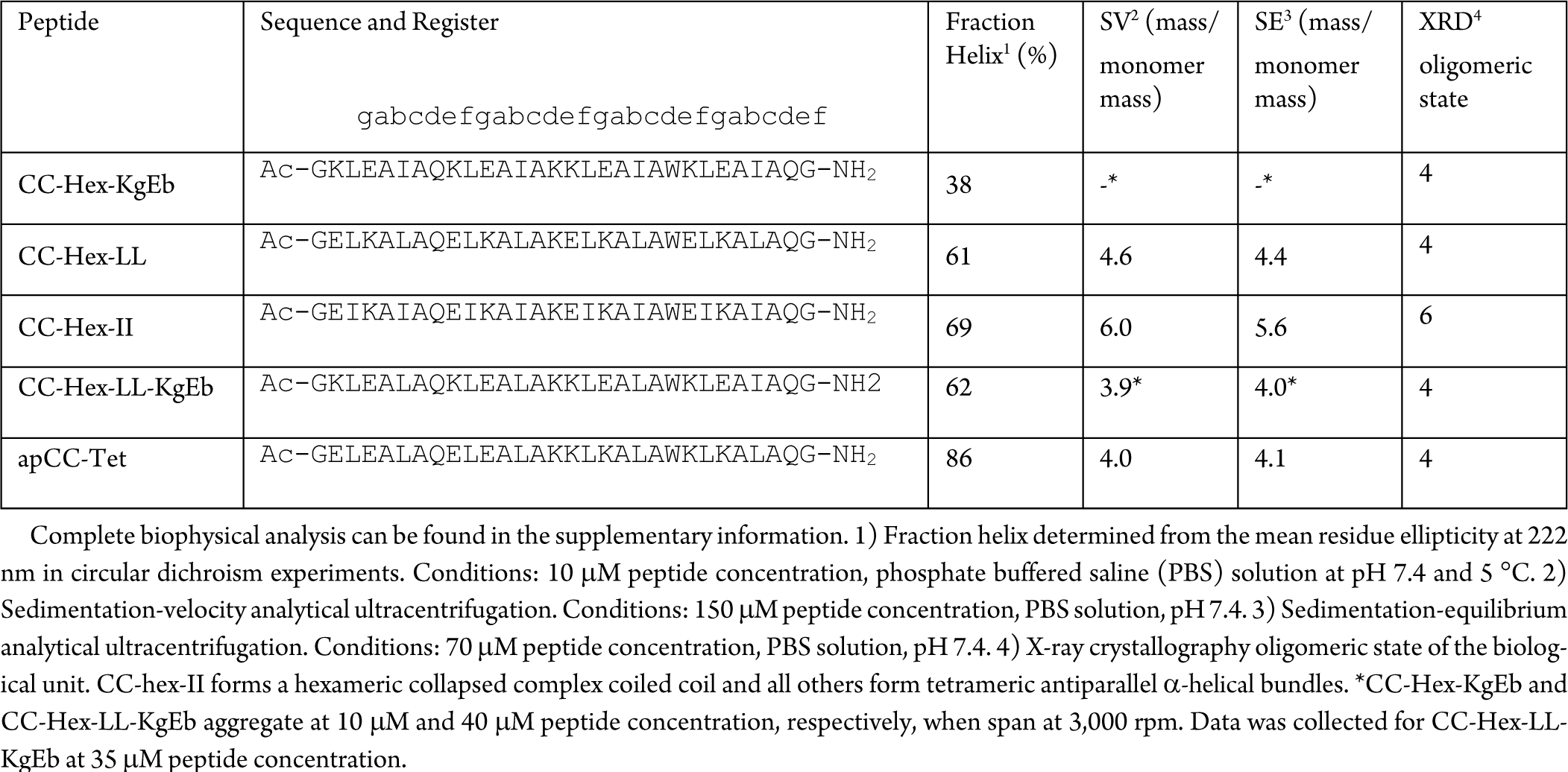
Sequences and summary of biophysical data for CC-Hex en bloc mutants and apCC-Tet.

First, we swapped the order of the charge residues at the *g* and *b* sites. In CC-Hex, the *g→f* repeat is ELKAIAx. Therefore, the swap had the sequence KLEAIAx, which we refer to as CC-Hex-KgEb. Surprisingly, this peptide was only soluble up to 40 μM concentration in phosphate buffered saline (PBS). It was 38% α helical but could not by analyzed by AUC due to aggregation. Nevertheless, we obtained an X-ray crystal structure of a variant that revealed an antiparallel tetramer (Figure 4A). Unlike the L24 variants, this does not have a shift between antiparallel interfaces and, consequently, it is a blunt-ended tetramer.

**Figure 4.**
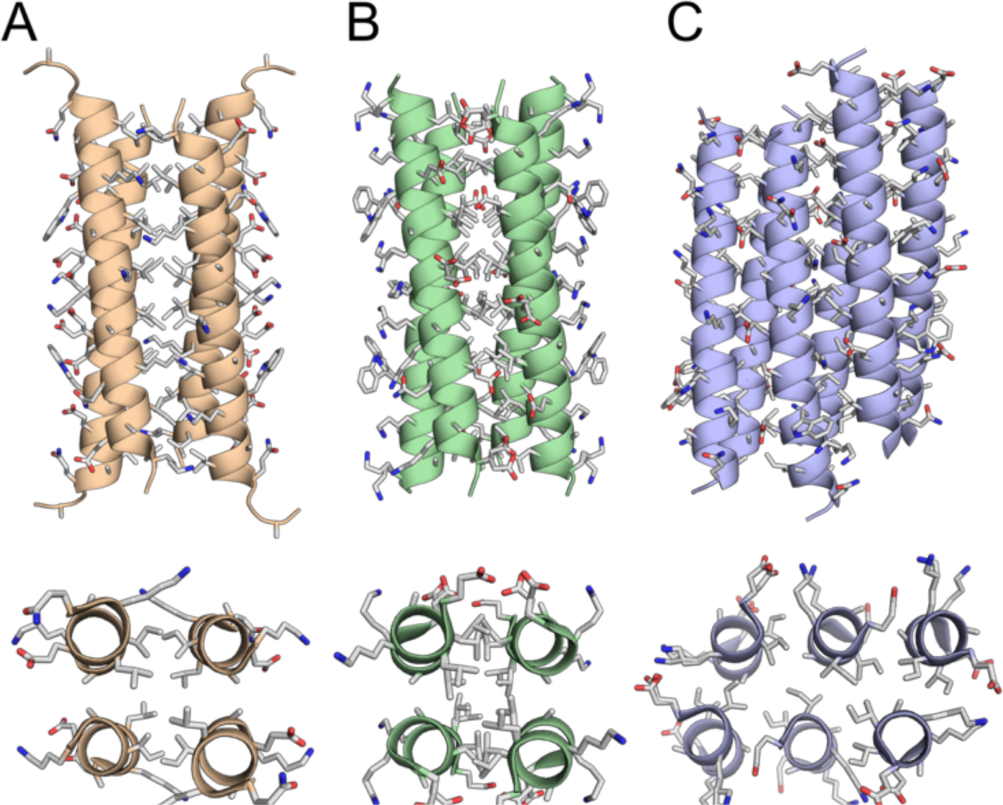
X-ray crystal structures of CC-Hex *en bloc* mutants. (A, B) CC-Hex-KgEb (orange) and CC-Hex-LL (green) form antiparallel tetramers. (C) CC-Hex-II (blue) forms a collapsed parallel hexamer.

Thus, swapping the potential salt-bridge positions disrupts the coiled-coil assembly. Analysis of KIH interactions in the CC-Hex structure reveals Glu at *g* as knob residues, with the ethyl unit (Cα and C*γ*) packing into the interface and the carboxylic acid presented at the surface. By contrast, the Lys side chains at *b* extend fully into solution. Given this, it is surprising that the swap is not tolerated: it is reasonable to expect that the methylene groups of Lys should pack at *g*, and the Glu at *b* should be solvent accessible. We speculate that the entropic cost of reducing the conformational freedom of Lys side chains at *g* disfavors hexamer formation.

Next, we turned to the *en bloc* hydrophobic mutations at *a* and *d* positions of CC-Hex. One of these mutants has already been reported: swapping the Leu at *a* and Ile at *d* results in the formation of a slipped hexameric barrel, CC-Hex-IL.^68^ For the study presented herein, we made the two other permutants with either all Ile or all Leu at both *a* and *d, i.e.*, CC-Hex-II and CC-Hex-LL, respectively. CC-Hex-II was α helical and completely unfolded upon heating, whereas CC-Hex-LL was highly α helical and hyperthermal stable (Figures S4.7). CC-Hex-LL showed the start of a thermal unfolding transition at ∼ 75 °C and, on cooling, signs of thermal annealing. AUC experiments indicated that CC-Hex-II was hexameric in solution, whereas CC-Hex-LL formed species of lower mass (Table 2).

X-ray crystal structures for both CC-Hex-II and CC-Hex-LL were determined (Figure 4B,C). The former revealed a collapsed hexameric coiled coil. This is a complex coiled coil with multiple unique helical environments in the homomeric assembly. Recently, we have described similar homomeric structures in which symmetry is broken.^69^ Albeit in a different *e/g* background, we argue that *β*-branched residues at the *a/d* sites promote barrel structures.^69^ Clearly, this is not the case for the new peptides described here: the introduction of additional Ile residues to CC-Hex to give CC-Hex-II results in a structure with a consolidated core. This demonstrates that residues peripheral to core sites also contribute to the final structure that is adopted. Previously, we solved an alternative structure for a variant of the CC-Hex-II peptide, which is a parallel tetramer (PDB accession code: 4H7R). The crystal structure for CC-Hex-LL also revealed an antiparallel tetramer. As with CC-Hex-KgEb, the peptide chains are fully helical. However, the backbones of these two structures could not be aligned (Figure 4A,B): the narrow Ala@*e* interface of CC-Hex-LL is longitudinally offset to a lesser extent than in CC-Hex-KgEb resulting in more-flushed termini.

These results demonstrate further the complexity of self-association landscape of CC-Hex-based peptides. And that alternative states can be accessed by changes in solution conditions and/or small changes to the sequence. Nonetheless, they also suggest ways in which structures within this landscape can be targeted, which we illustrate next.

### Designing apCC-Tet, an optimized antiparallel coiled-coil tetramer

From the above, we chose CC-Hex-LL as the best starting point for the core in a rational design of a fully folded, stable antiparallel homotetramer. In addition, we swapped the charged residues at *g* and *b* positions, as this also favors antiparallel tetramer over hexamer. This gave CC-Hex-LL-KgEb (Table 2). Compared with CC-Hex-LL, CC-Hex-LL-KgEb gave a sharp single peak in the c(s) distribution in sedimentation velocity AUC. Sedimentation-equilibrium data and an X-ray crystal structure confirmed CC-Hex-LL-KgEb as a tetramer (Figure 5). However, like CC-Hex-KgEb, the new variant aggregated in PBS above 40 μM concentration. Aggregation of the tetramers could occur along the wide interface of these oblate structures; relative to the narrow-interface, the wide interfaces present a greater exposed hydrophobic surface. It is possible that peripheral lysine residues screen this interface less well than glutamate. Thus, whilst swapping the potential salt bridges help specify the tetrameric form relative to the hexamer this is at the cost of reduced solubility. Therefore, we sought an alternate and optimized pattern of charge on the exterior of the tetramer.

**Figure 5.**
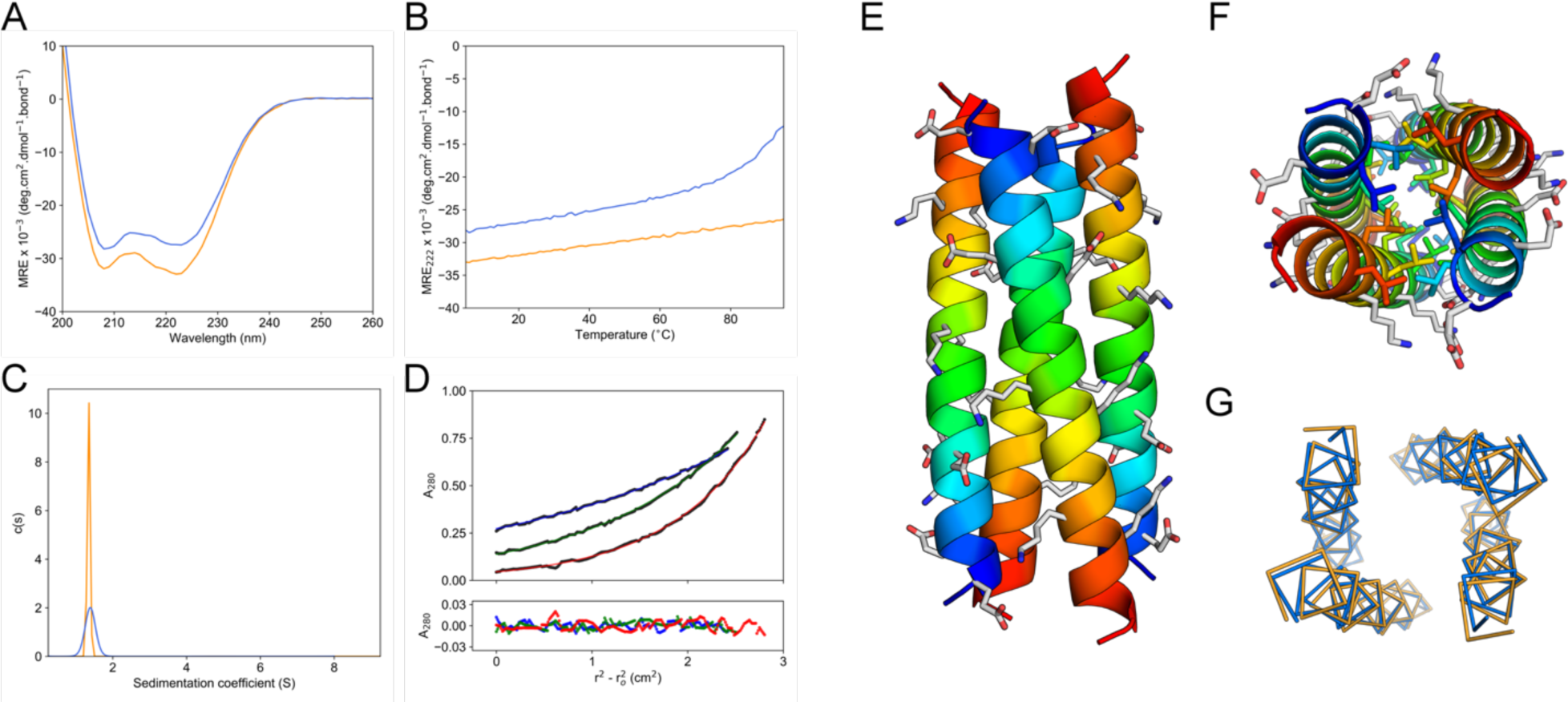
(A) Circular dichroism (CD) spectra at 5 °C and (B) thermal unfolding of apCC-Tet (orange) and CC-Hex-LL (blue). Thermal unfolding of CC-Hex-LL was performed after thermally annealing the sample. Conditions: 10 µM peptide, PBS. (C) Sedimentation-velocity AUC of apCC-Tet (orange) and CC-Hex-LL (blue). Fits return masses of 4.0 and 4.6, respectively. Conditions: 150 µM peptide, PBS. (D) Sedimentation-equilibrium AUC of apCC-Tet (blue, 24k rpm; green, 32k rpm; and red, 40k rpm). Fitting to the experimental data returned a weight of 4.1 x monomer mass. Conditions: 70 µM peptide concentration, PBS. (E,F) X-ray crystal structure of apCC-Tet colored from *N* (blue) to *C* terminus (red). Side chains of the *g* and *b* residues and core Leu residues are shown as sticks. (G) Backbone comparison of apCC-Tet (orange) and CC-Hex-LL (blue).

There are several ways to arrange Lys and Glu residues at *g* and *b* positions to disfavor parallel association of helices but favor the antiparallel alignment.^70–74^ For example, Glu could be placed at *g* and *b* of heptads 1 and 2, with Lys at these sites in heptads 3 and 4. This arrangement of Glu near to the *N* terminus and Lys near the *C* terminus is known to improve α-helical stability.^75^ Retrospectively, we named this charge pattern in the CC-Hex-LL background apCC-Tet.

apCC-Tet was highly α helical at 5 °C with a fraction helicity of 86%, an increase of 25% over CC-Hex-LL (Figure 5A). apCC-Tet was thermally stable up to 95 °C, whereas CC-Hex-LL showed the start of an unfolding transition at ∼ 75 °C (Figure 5B). This improvement in solution-phase properties of apCC-Tet was also evident in AUC experiments (Figure 5C,D). In sedimentation-velocity AUC, apCC-Tet gave a much sharper peak. apCC-Tet gave a single discrete species with a weight of 4.0 and 4.1 x monomer mass by sedimentation velocity and sedimentation equilibrium, respectively; whereas, CC-Hex-LL returned non-integer oligomeric states of 4.6 and 4.4, respectively. Crucially, apCC-Tet did not show any signs of aggregation at 150 μM peptide concentration during these experiments. Collectively, these data show that the arrangement of charged residues that should favor antiparallel helices also has improves the folding and thermal stability of apCC-Tet.

The X-ray crystal structure for apCC-Tet confirmed an antiparallel tetramer (Figure 5D-F). The helix geometry and interfaces are similar to CC-Hex-LL with backbone and all-atom RMSDs of 0.56 Å and 1.58 Å between the two structures. However, only 7 of the possible 16 pairs of Glu-Lys contacts in the apCC-Tet structure are within 4 Å to form interhelical salt bridges and the side chains are diffuse (*e.g.* the average temperature factor for the Lys N*ζ* is 56.9 Å^2^ in comparison the all-atom average is 40.5 Å^2^). Therefore, despite welcome improvements in the solution-phase behavior, the X-ray structure reveals that these are not due to altered backbone arrangements or core packing, nor are they from salt-bridge formation. We posit that the changes allow electrostatic steering in helix association and avoid uniformly charged surfaces on apCC-Tet.

To summarize this section, we have achieved the rational design of an antiparallel homomeric coiled-coil tetramer. In the register of the heptad repeat, this has a hydrophobic core of *a* = *d* = Leu plus *e* = Ala, with flanking Glu-Lys pairs at *g:g* and *b:b* sites. This sequence pattern directs the assembly of a well-defined, discrete and hyperstable antiparallel tetramer with an up-down-up-down topology and oblate cross-section. This adds to a growing basis set of *de novo* CC modules for protein design and synthetic biology.

## CONCLUSION

CC-Hex was the first reported X-ray crystal structure of a *de novo* α-helical barrel.^60^ These barrels are a growing class of both natural and designed proteins that are robust and have potential in protein engineering and synthetic biology.^55,62–64,76,77^ Nonetheless, CC-Hex was discovered serendipitously: it was a variant of a *de novo* designed parallel coiled-coil tetramer, CC-Tet.^59^ Herein, we have described a series of point and *en bloc* changes to the CC-Hex sequence. These reveal that this background is somewhat plastic with a variety of coiledcoil states accessible from it, including parallel and antiparallel tetramers, and open barrel and collapsed hexamers. Moreover, the structural plasticity is evident even for some of the charge-bearing point variants, which can be switched from the lower to higher oligomeric states by neutralizing the charge of the introduced side chains. These empirical explorations and observations have allowed us to design *de novo* a new antiparallel homotetrameric coiled coil, apCC-Tet. A combination of the established positive and negative design rules resulted in a robust and fully characterized structure.

The work presented here highlights further the degrees of freedom available to self-associating peptide systems,^56,57,69^ and the associated challenge of designing specific oligomer states and topologies *de novo*.^12,55,78–80^ This problem is particularly acute for less-well-defined peptide-peptide interfaces where many near-isoenergetic states are accessible.^12^

The energy landscape for α-helical coiled-coil assemblies may well be more navigable than the general case of helical bundles. This is for three interrelated reasons: First, the hallmark knobs-into-holes packing of coiled coils dramatically reduces the number of helical arrangements possible. As a result, it is relatively straightforward to enumerate many coiled-coil backbones parametrically and/or computationally.^81–86^ Second, and directly related to this structural constraint, coiled-coil sequences have relatively simple repeat patterns of hydrophobic and polar residues. Third, and as a consequence of the first two points, only certain residues appear to be tolerated in the helix-helix interfaces of coiled coils.^78–80^ Thus, as coiled coils are low-complexity sequences, the protein-design problem is more sharply defined for these structures and sequences compared with less-regular structures.

That said, the coiled-coil energy landscape is still complex with multiple oligomer states, parallel/antiparallel/mixed arrangements of helices, and homo- and heterotypic assemblies all possible.^54, 87^ Nonetheless, considerable progress has been made to discern sequence-to-structure relationships and computational methods for coiled-coil design. These have led to robust rational and/or computational designs for parallel dimers through heptamers.^59,60,62,76^ The work presented herein adds to this effort: it illustrates how alternate states can be distinguished; and it provides guidelines for accessing *de novo* antiparallel structures, which have been less explored than parallel coiled coils.^71–73,88^

More subtly, certain polar residues are tolerated at the otherwise hydrophobic coiled-coil interfaces, and these influence partner and oligomer-state selection.^78,89,90^ This concept has recently been revisited.^91, 92^ Here, we add to this showing that buried charged residues disfavor high-oligomer states for alternate antiparallel tetramers in the CC-Hex background. Furthermore, for certain sequences the two states—parallel hexamer and antiparallel tetramer—are sufficiently close in energy to effect switches between them simply by changing the pH of the solution. This presents exciting prospects for developing sequences to switch gross structural state in response to facile perturbations.^70,93–97^

Coiled-coil peptides that show multiple-defined states have been described by others. Lizatović *et al.* design a pH-responsive sequence that switches between a pentameric bundle and parallel hexameric bundle.^93^ Grigoryan *et al.* present the design of carbon-nanotube solubilizing peptides, which wrap around the surface of the nanotubes. In isolation, one of these peptides forms a tetramer, and another forms a dimer/hexamer mixture.^66^ Others are also discovering that seemingly benign sequence alterations can cause gross structural changes in self-associating coiled coils. Slovic *et al.* describe the redesign of the membrane-spanning peptide phospholambin to make it water soluble. Solution-phase biophysics shows that the full-length sequence remains pentameric. However, a truncated peptide (residues 21-52) is best modelled as a tetramer/pentamer equilibrium in AUC and a structure reveals an offset antiparallel tetramer.^98, 99^ Finally, Spencer *et al.* have made variants of CC-Hex with phenylalanine in the core of CC-Hex that form a collapsed antiparallel hexameric bundle.^100, 101^

The specification of orientation of peptides in self-associating systems is a standing challenge in protein design. Antiparallel coiled coils remain a challenging design target with fewer examples of successful designs compared to parallel structures. We have delivered a robust *de novo* antiparallel tetramer that has been thoroughly characterized. This provides another designed module or component adding to the growing basis set of *de novo* coiled coils with potential applications in the design and assembly of protein origami,^102^ peptide nanotubes,^68^ protein colocalization,^103, 104^ and generally as tectons for generating complex self-assembling systems.^105, 106^

## ASSOCIATED CONTENT

### Supporting Information

Supporting Figures and Tables (PDF)

### Crystal-Structure Files

The crystal structures are all available from the Protein Data Bank using the following accession codes: CC-Hex-L24D (6Q5H), CC-Hex-L24E_tet (6Q5I), CC-Hex-L24E_hex (6Q5J), CC-Hex-L24K (6Q5K), CC-Hex-L24H (6Q5L), CC-Hex-L24DAB (6Q5M), CC-Hex-L24Nle (6Q5N), CC-Hex-LL (6Q5O), CC-Hex-II (6Q5P), CC-Hex-KgEb_var (6Q5Q), CC-Hex-LL-KgEb (6Q5R), apCC-Tet (6Q5S).

## AUTHOR INFORMATION

### Corresponding Author

*

### Author Contributions

G.G.R., C.W.W., J.L.B., A.R.T. and D.N.W. conceived the project and designed the experiments. G.G.R., C.W.W., J.L.B. and A.J.B. synthesized the peptides and conducted the solution-phase biophysics. G.G.R., C.W.W., J.L.B., N.R.Z., A.J.B. and R.L.B. determined the peptide X-ray crystal structures. G.G.R., C.W.W. and D.N.W. wrote the paper. All authors have read and contributed to the preparation of the manuscript.

### Notes

The authors declare no competing financial interest.

## ACKNOWLEDGMENT

G.G.R., C.W.W., J.L.B., A.R.T., A.J.B. and DNW are supported by a European Research Council Advanced Grant to DNW (340764). G.G.R. and A.J.B. thank the Bristol Chemical Synthesis Centre for Doctoral Training funded by the Engineering and Physical Sciences Research Council (EP/G036764/1). C.W.W. and D.N.W thank the Biotechnology and Biological Sciences Research Council for funding the South West Biosciences Doctoral Training Partnership (BB/J014400/1) and a responsive-mode grant (BB/R00661X/1). We thank the University of Bristol School of Chemistry Mass Spectrometry Facility for access to the EPSRC-funded Bruker Ultraflex MALDI-TOF/TOF instrument (EP/K03927X/1). We thank the MX group at Diamond Light Source for their support. D.N.W. holds a Royal Society Wolfson Research Merit Award (WM140008). D.N.W. would like to dedicate this paper to Professor Ronald T Raines on the occasion of his 60th birthday.

---

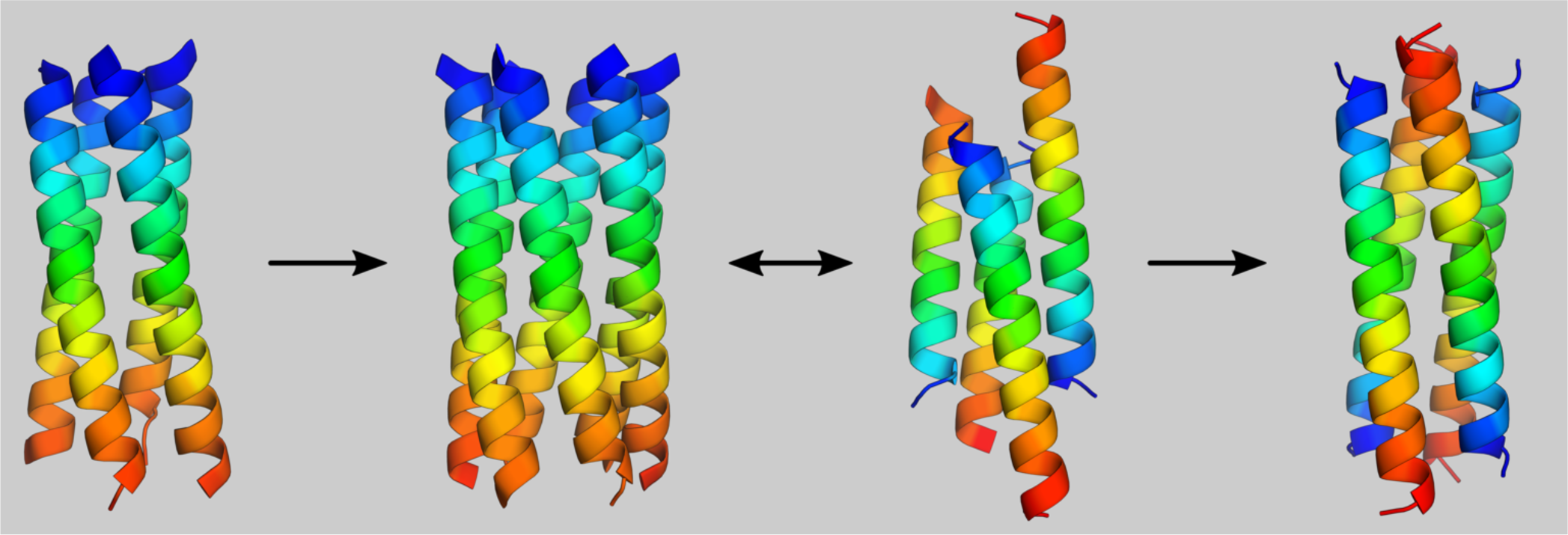

## REFERENCES

(1) Vos, A. de; Ultsch, M.; Kossiakoff, A. A. Human Growth Hormone and Extracellular Domain of Its Receptor: Crystal Structure of the Complex. Science 1992, 255 (5042), 306–312. https://doi.org/10.1126/science.1549776.

(2) Powers, R.; Garrett, D. S.; March, C. J.; Frieden, E. A.; Gronenborn, A. M.; Clore, G. M. The High-Resolution, Three-Dimensional Solution Structure of Human Interleukin-4 Determined by Multidimensional Heteronuclear Magnetic Resonance Spectroscopy. Biochemistry 1993, 32 (26), 6744–6762. https://doi.org/10.1021/bi00077a030.

(3) Lederer, F.; Glatigny, A.; Bethge, P. H.; Bellamy, H. D.; Mathews, F. S. Improvement of the 2.5 Å Resolution Model of Cytochrome B562 by Redetermining the Primary Structure and Using Molecular Graphics. Journal of Molecular Biology 1981, 148 (4), 427–448. https://doi.org/10.1016/0022-2836(81)90185-6.

(4) Klabunde, T.; Eicken, C.; Sacchettini, J. C.; Krebs, B. Crystal Structure of a Plant Catechol Oxidase Containing a Di-copper Center. Nature Structural & Molecular Biology 1998, 5 (12), 1084–1090. https://doi.org/10.1038/4193.

(5) Lawson, D. M.; Artymiuk, P. J.; Yewdall, S. J.; Smith, J. M. A.; Livingstone, J. C.; Treffry, A.; Luzzago, A.; Levi, S.; Arosio, P.; Cesareni, G.; et al. Solving the Structure of Human H Ferritin by Genetically Engineering Intermolecular Crystal Contacts. Nature 1991, 349 (6309), 541–544. https://doi.org/10.1038/349541a0.

(6) Friedman, A. M.; Fischmann, T. O.; Steitz, T. A. Crystal Structure of Lac Repressor Core Tetramer and Its Implications for DNA Looping. Science 1995, 268 (5218), 1721–1727. https://doi.org/10.1126/science.7792597.

(7) Ferré-D’Amaré, A. R.; Prendergast, G. C.; Ziff, E. B.; Burley, S. K. Recognition by Max of Its Cognate DNA through a Dimeric b/HLH/Z Domain. Nature 1993, 363 (6424), 38–45. https://doi.org/10.1038/363038a0.

(8) Sutton, R. B.; Fasshauer, D.; Jahn, R.; Brunger, A. T. Crystal Structure of a SNARE Complex Involved in Synaptic Exocytosis at 2.4 Å Resolution. Nature 1998, 395 (6700), 347–353. https://doi.org/10.1038/26412.

(9) Wynne, S. A.; Crowther, R. A.; Leslie, A. G. W. The Crystal Structure of the Human Hepatitis B Virus Capsid. Molecular Cell 1999, 3 (6), 771–780. https://doi.org/10.1016/S1097-2765(01)80009-5.

(10) Hulko, M.; Berndt, F.; Gruber, M.; Linder, J. U.; Truffault, V.; Schultz, A.; Martin, J.; Schultz, J. E.; Lupas, A. N.; Coles, M. The HAMP Domain Structure Implies Helix Rotation in Transmembrane Signaling. Cell 2006, 126 (5), 929–940. https://doi.org/10.1016/j.cell.2006.06.058.

(11) Thomaston, J. L.; Alfonso-Prieto, M.; Woldeyes, R. A.; Fraser, J. S.; Klein, M. L.; Fiorin, G.; DeGrado, W. F. High-Resolution Structures of the M2 Channel from Influenza A Virus Reveal Dynamic Pathways for Proton Stabilization and Transduction. PNAS 2015, 112 (46), 14260–14265. https://doi.org/10.1073/pnas.1518493112.

(12) Hill, R. B.; Raleigh, D. P.; Lombardi, A.; DeGrado, W. F. De Novo Design of Helical Bundles as Models for Understanding Protein Folding and Function. Acc. Chem. Res. 2000, 33 (11), 745–754. https://doi.org/10.1021/ar970004h.

(13) Woolfson, D. N. Core-Directed Protein Design. Current Opinion in Structural Biology 2001, 11 (4), 464–471. https://doi.org/10.1016/S0959-440X(00)00234-7.

(14) Hecht, M. H.; Das, A.; Go, A.; Bradley, L. H.; Wei, Y. De Novo Proteins from Designed Combinatorial Libraries. Protein Science 2004, 13 (7), 1711–1723. https://doi.org/10.1110/ps.04690804.

(15) Horne, W. S.; Gellman, S. H. Foldamers with Heterogeneous Backbones. Acc. Chem. Res. 2008, 41 (10), 1399–1408. https://doi.org/10.1021/ar800009n.

(16) Chino, M.; Maglio, O.; Nastri, F.; Pavone, V.; DeGrado, W. F.; Lombardi, A. Artificial Diiron Enzymes with a De Novo Designed Four-Helix Bundle Structure. European Journal of Inorganic Chemistry 2015, 2015 (21), 3371–3390. https://doi.org/10.1002/ejic.201500470.

(17) Regan, L.; Caballero, D.; Hinrichsen, M. R.; Virrueta, A.; Williams, D. M.; O’Hern, C. S. Protein Design: Past, Present, and Future. Peptide Science 2015, 104 (4), 334–350. https://doi.org/10.1002/bip.22639.

(18) Grayson, K. J.; Anderson, J. L. R. Designed for Life: Biocompatible de Novo Designed Proteins and Components. Journal of The Royal Society Interface 2018, 15 (145), 20180472. https://doi.org/10.1098/rsif.2018.0472.

(19) Regan, L.; DeGrado, W. F. Characterization of a Helical Protein Designed from First Principles. Science 1988, 241 (4868), 976–978. https://doi.org/10.1126/science.3043666.

(20) Argos, P.; Rossmann, M. G.; Johnson, J. E. A Four-Helical Super-Secondary Structure. Biochemical and Biophysical Research Communications 1977, 75 (1), 83–86. https://doi.org/10.1016/0006-291X(77)91292-X.

(21) Crick, F. H. The Packing of α-Helices: Simple Coiled-Coils. Acta crystallographica 1953, 6 (8–9), 689–697.

(22) Weber, P. C.; Salemme, F. R. Structural and Functional Diversity in 4-α-Helical Proteins. Nature 1980, 287 (5777), 82–84. https://doi.org/10.1038/287082a0.

(23) Presnell, S. R.; Cohen, F. E. Topological Distribution of Four-Alpha-Helix Bundles. PNAS 1989, 86 (17), 6592–6596. https://doi.org/10.1073/pnas.86.17.6592.

(24) Harris, N. L.; Presnell, S. R.; Cohen, F. E. Four Helix Bundle Diversity in Globular Proteins. Journal of Molecular Biology 1994, 236 (5), 1356–1368. https://doi.org/10.1016/0022-2836(94)90063-9.

(25) Murzin, A. G.; Finkelstein, A. V. General Architecture of the α-Helical Globule. Journal of Molecular Biology 1988, 204 (3), 749–769. https://doi.org/10.1016/0022-2836(88)90366-X.

(26) Cohen, C.; Parry, D. A. D. α-Helical Coiled Coils and Bundles: How to Design an α-Helical Protein. Proteins: Structure, Function, and Bioinformatics 1990, 7 (1), 1–15. https://doi.org/10.1002/prot.340070102.

(27) Bryson, J. W.; Betz, S. F.; Lu, H. S.; Suich, D. J.; Zhou, H. X.; O’Neil, K. T.; DeGrado, W. F. Protein Design: A Hierarchic Approach. Science 1995, 270 (5238), 935–941. https://doi.org/10.1126/science.270.5238.935.

(28) Chothia, C.; Levitt, M.; Richardson, D. Structure of Proteins: Packing of Alpha-Helices and Pleated Sheets. Proceedings of the National Academy of Sciences 1977, 74 (10), 4130–4134.

(29) Chothia, C.; Levitt, M.; Richardson, D. Helix to Helix Packing in Proteins. Journal of molecular biology 1981, 145 (1), 215–250.

(30) DeGrado, W. F.; Wasserman, Z. R.; Lear, J. D. Protein Design, a Minimalist Approach. Science 1989, 243 (4891), 622–628. https://doi.org/10.1126/science.2464850.

(31) Ho, S. P.; DeGrado, W. F. Design of a 4-Helix Bundle Protein: Synthesis of Peptides Which Self-Associate into a Helical Protein. J. Am. Chem. Soc. 1987, 109 (22), 6751–6758. https://doi.org/10.1021/ja00256a032.

(32) Hill, C. P.; Anderson, D. H.; Wesson, L.; DeGrado, W. F.; Eisenberg, D. Crystal Structure of Alpha 1: Implications for Protein Design. Science 1990, 249 (4968), 543–546. https://doi.org/10.1126/science.2382133.

(33) Handel, T.; DeGrado, W. F. De Novo Design of a Zn2+-Binding Protein. J. Am. Chem. Soc. 1990, 112 (18), 6710–6711. https://doi.org/10.1021/ja00174a039.

(34) Raleigh, D. P.; DeGrado, W. F. A de Novo Designed Protein Shows a Thermally Induced Transition from a Native to a Molten Globule-like State. J. Am. Chem. Soc. 1992, 114 (25), 10079–10081. https://doi.org/10.1021/ja00051a061.

(35) Raleigh, D. P.; Betz, S. F.; DeGrado, W. F. A de Novo Designed Protein Mimics the Native State of Natural Proteins. J. Am. Chem. Soc. 1995, 117 (28), 7558–7559. https://doi.org/10.1021/ja00133a035.

(36) Hill, R. B.; DeGrado, W. F. Solution Structure of Α2D, a Nativelike de Novo Designed Protein. J. Am. Chem. Soc. 1998, 120 (6), 1138–1145. https://doi.org/10.1021/ja9733649.

(37) Joh, N. H.; Wang, T.; Bhate, M. P.; Acharya, R.; Wu, Y.; Grabe, M.; Hong, M.; Grigoryan, G.; DeGrado, W. F. De Novo Design of a Transmembrane Zn2+-Transporting Four-Helix Bundle. Science 2014, 346 (6216), 1520–1524. https://doi.org/10.1126/science.1261172.

(38) Polizzi, N. F.; Wu, Y.; Lemmin, T.; Maxwell, A. M.; Zhang, S.-Q.; Rawson, J.; Beratan, D. N.; Therien, M. J.; DeGrado, W. F. De Novo Design of a Hyperstable Non-Natural Protein–Ligand Complex with Sub-Å Accuracy. Nature Chemistry 2017, 9 (12), 1157–1164. https://doi.org/10.1038/nchem.2846.

(39) Gibney, B. R.; Dutton, P. L. De Novo Design and Synthesis of Heme Proteins. In Advances in Inorganic Chemistry; Academic Press, 2000; Vol. 51, pp 409–456. https://doi.org/10.1016/S0898-8838(00)51008-3.

(40) Koder, R. L.; Dutton, P. L. Intelligent Design: The de Novo Engineering of Proteins with Specified Functions. Dalton Trans. 2006, 0 (25), 3045–3051. https://doi.org/10.1039/B514972J.

(41) Koder, R. L.; Anderson, J. L. R.; Solomon, L. A.; Reddy, K. S.; Moser, C. C.; Dutton, P. L. Design and Engineering of an O2 Transport Protein. Nature 2009, 458 (7236), 305–309. https://doi.org/10.1038/nature07841.

(42) Farid, T. A.; Kodali, G.; Solomon, L. A.; Lichtenstein, B. R.; Sheehan, M. M.; Fry, B. A.; Bialas, C.; Ennist, N. M.; Siedlecki, J. A.; Zhao, Z.; et al. Elementary Tetrahelical Protein Design for Diverse Oxidoreductase Functions. Nature Chemical Biology 2013, 9 (12), 826–833. https://doi.org/10.1038/nchem-bio.1362.

(43) Kodali, G.; A. Mancini, J.; A. Solomon, L.; V. Episova, T.; Roach, N.; J. Hobbs, C.; Wagner, P.; A. Mass, O.; Aravindu, K.; E. Barnsley, J.; et al. Design and Engineering of Water-Soluble Light-Harvesting Protein Maquettes. Chemical Science 2017, 8 (1), 316–324. https://doi.org/10.1039/C6SC02417C.

(44) Watkins, D. W.; Jenkins, J. M. X.; Grayson, K. J.; Wood, N.; Steventon, J. W.; Vay, K. K. L.; Goodwin, M. I.; Mullen, A. S.; Bailey, H. J.; Crump, M. P.; et al. Construction and in Vivo Assembly of a Catalytically Proficient and Hyperthermostable de Novo Enzyme. Nature Communications 2017, 8 (1), 358. https://doi.org/10.1038/s41467-017-00541-4.

(45) Kamtekar, S.; Schiffer, J. M.; Xiong, H.; Babik, J. M.; Hecht, M. H. Protein Design by Binary Patterning of Polar and Nonpolar Amino Acids. Science 1993, 262 (5140), 1680–1685. https://doi.org/10.1126/science.8259512.

(46) Wei, Y.; Liu, T.; Sazinsky, S. L.; Moffet, D. A.; Pelczer, I.; Hecht, M. H. Stably Folded de Novo Proteins from a Designed Combinatorial Library. Protein Science 2003, 12 (1), 92–102. https://doi.org/10.1110/ps.0228003.

(47) Wei, Y.; Kim, S.; Fela, D.; Baum, J.; Hecht, M. H. Solution Structure of a de Novo Protein from a Designed Combinatorial Library. PNAS 2003, 100 (23), 13270–13273. https://doi.org/10.1073/pnas.1835644100.

(48) Hecht, M. H.; Vogel, K. M.; Spiro, T. G.; Rojas, N. R. L.; Kamtekar, S.; Simons, C. T.; Mclean, J. E.; Farid, R. S. De Novo Heme Proteins from Designed Combinatorial Libraries. Protein Science 1997, 6 (12), 2512–2524. https://doi.org/10.1002/pro.5560061204.

(49) Fisher, M. A.; McKinley, K. L.; Bradley, L. H.; Viola, S. R.; Hecht, M. H. De Novo Designed Proteins from a Library of Artificial Sequences Function in Escherichia Coli and Enable Cell Growth. PLOS ONE 2011, 6 (1), e15364. https://doi.org/10.1371/journal.pone.0015364.

(50) Murphy, G. S.; Greisman, J. B.; Hecht, M. H. De Novo Proteins with Life-Sustaining Functions Are Structurally Dynamic. Journal of Molecular Biology 2016, 428 (2, Part A), 399–411. https://doi.org/10.1016/j.jmb.2015.12.008.

(51) Cherny, I.; Korolev, M.; Koehler, A. N.; Hecht, M. H. Proteins from an Unevolved Library of de Novo Designed Sequences Bind a Range of Small Molecules. ACS Synth. Biol. 2012, 1 (4), 130–138. https://doi.org/10.1021/sb200018e.

(52) Hoegler, K. J.; Hecht, M. H. A de Novo Protein Confers Copper Resistance in E Scherichia Coli. Protein Sci 2016, 25 (7), 1249–1259. https://doi.org/10.1002/pro.2871.

(53) Lombardi, A.; Summa, C. M.; Geremia, S.; Randaccio, L.; Pavone, V.; DeGrado, W. F. Retrostructural Analysis of Metalloproteins: Application to the Design of a Minimal Model for Diiron Proteins. PNAS 2000, 97 (12), 6298–6305. https://doi.org/10.1073/pnas.97.12.6298.

(54) Lupas, A. N.; Bassler, J.; Dunin-Horkawicz, S. The Structure and Topology of α-Helical Coiled Coils. In Fibrous Proteins: Structures and Mechanisms; Parry, D. A. D., Squire, J. M., Eds.; Subcellular Biochemistry; Springer International Publishing: Cham, 2017; pp 95–129. https://doi.org/10.1007/978-3-319-49674-0_4.

(55) Woolfson, D. N. Coiled-Coil Design: Updated and Upgraded. In Fibrous Proteins: Structures and Mechanisms; Parry, D. A. D., Squire, J. M., Eds.; Subcellular Biochemistry; Springer International Publishing: Cham, 2017; pp 35–61. https://doi.org/10.1007/978-3-319-49674-0_2.

(56) Harbury, P. B.; Zhang, T.; Kim, P. S.; Alber, T. A Switch between Two-, Three-, and Four-Stranded Coiled Coils in GCN4 Leucine Zipper Mutants. Science 1993, 262 (5138), 1401–1407.

(57) Harbury, P. B.; Kim, P. S.; Alber, T. Crystal Structure of an Isoleucine-Zipper Trimer. Nature 1994, 371 (6492), 80.

(58) Woolfson, D. N. The Design of Coiled-Coil Structures and Assemblies. In Advances in Protein Chemistry; Fibrous Proteins: Coiled-Coils, Collagen and Elastomers; Academic Press, 2005; Vol. 70, pp 79–112. https://doi.org/10.1016/S0065-3233(05)70004-8.

(59) Fletcher, J. M.; Boyle, A. L.; Bruning, M.; Bartlett, G. J.; Vincent, T. L.; Zaccai, N. R.; Armstrong, C. T.; Bromley, E. H.; Booth, P. J.; Brady, R. L. A Basis Set of de Novo Coiled-Coil Peptide Oligomers for Rational Protein Design and Synthetic Biology. ACS synthetic biology 2012, 1 (6), 240–250.

(60) Zaccai, N. R.; Chi, B.; Thomson, A. R.; Boyle, A. L.; Bartlett, G. J.; Bruning, M.; Linden, N.; Sessions, R. B.; Booth, P. J.; Brady, R. L.; et al. A de Novo Peptide Hexamer with a Mutable Channel. Nat Chem Biol 2011, 7 (12), 935–941. https://doi.org/10.1038/nchembio.692.

(61) Burton, A. J.; Thomas, F.; Agnew, C.; Hudson, K. L.; Halford, S. E.; Brady, R. L.; Woolfson, D. N. Accessibility, Reactivity, and Selectivity of Side Chains within a Channel of de Novo Peptide Assembly. J. Am. Chem. Soc. 2013, 135 (34), 12524–12527. https://doi.org/10.1021/ja4053027.

(62) Thomson, A. R.; Wood, C. W.; Burton, A. J.; Bartlett, G. J.; Sessions, R. B.; Brady, R. L.; Woolfson, D. N. Computational Design of Water-Soluble α-Helical Barrels. Science 2014, 346 (6208), 485–488. https://doi.org/10.1126/science.1257452.

(63) Burton, A. J.; Thomson, A. R.; Dawson, W. M.; Brady, R. L.; Woolfson, D. N. Installing Hydrolytic Activity into a Completely de Novo Protein Framework. Nat Chem 2016, 8 (9), 837–844. https://doi.org/10.1038/nchem.2555.

(64) Thomas, F.; Dawson, W. M.; Lang, E. J. M.; Burton, A. J.; Bartlett, G. J.; Rhys, G. G.; Mulholland, A. J.; Woolfson, D. N. De Novo-Designed α-Helical Barrels as Receptors for Small Molecules. ACS Synth. Biol. 2018, 7 (7), 1808–1816. https://doi.org/10.1021/acssynbio.8b00225.

(65) Hass, M. A. S.; Mulder, F. A. A. Contemporary NMR Studies of Protein Electrostatics. Annu. Rev. Biophys. 2015, 44 (1), 53–75. https://doi.org/10.1146/annurev-biophys-083012-130351.

(66) Grigoryan, G.; Kim, Y. H.; Acharya, R.; Axelrod, K.; Jain, R. M.; Willis, L.; Drndic, M.; Kikkawa, J. M.; DeGrado, W. F. Computational Design of Virus-Like Protein Assemblies on Carbon Nanotube Surfaces. Science 2011, 332 (6033), 1071–1076. https://doi.org/10.1126/science.1198841.

(67) Thomas, F.; Boyle, A. L.; Burton, A. J.; Woolfson, D. N. A Set of de Novo Designed Parallel Heterodimeric Coiled Coils with Quantified Dissociation Constants in the Micromolar to Sub-Nanomolar Regime. J. Am. Chem. Soc. 2013, 135 (13), 5161–5166. https://doi.org/10.1021/ja312310g.

(68) Burgess, N. C.; Sharp, T. H.; Thomas, F.; Wood, C. W.; Thomson, A. R.; Zaccai, N. R.; Brady, R. L.; Serpell, L. C.; Woolfson, D. N. Modular Design of Self-Assembling Peptide-Based Nanotubes. J. Am. Chem. Soc. 2015, 137 (33), 10554–10562. https://doi.org/10.1021/jacs.5b03973.

(69) Rhys, G. G.; Wood, C. W.; Lang, E. J. M.; Mulholland, A. J.; Brady, R. L.; Thomson, A. R.; Woolfson, D. N. Maintaining and Breaking Symmetry in Homomeric Coiled-Coil Assemblies. Nature Communications 2018, 9 (1), 4132. https://doi.org/10.1038/s41467-018-06391-y.

(70) Pandya, M. J.; Cerasoli, E.; Joseph, A.; Stoneman, R. G.; Waite, E.; Woolfson, D. N. Sequence and Structural Duality:?Designing Peptides to Adopt Two Stable Conformations. J. Am. Chem. Soc. 2004, 126 (51), 17016–17024. https://doi.org/10.1021/ja045568c.

(71) Gurnon, D. G.; Whitaker, J. A.; Oakley, M. G. Design and Characterization of a Homodimeric Antiparallel Coiled Coil. J. Am. Chem. Soc. 2003, 125 (25), 7518–7519. https://doi.org/10.1021/ja0357590.

(72) McClain, D. L.; Woods, H. L.; Oakley, M. G. Design and Characterization of a Heterodimeric Coiled Coil That Forms Exclusively with an Antiparallel Relative Helix Orientation. J. Am. Chem. Soc. 2001, 123 (13), 3151–3152. https://doi.org/10.1021/ja004099l.

(73) McClain, D. L.; Binfet, J. P.; Oakley, M. G. Evaluation of the Energetic Contribution of Interhelical Coulombic Interactions for Coiled Coil Helix Orientation Specificity. J Mol Biol 2001, 313 (2), 371–383. https://doi.org/10.1006/jmbi.2001.5044.

(74) Kohn, W. D.; Kay, C. M.; Hodges, R. S. Protein Destabilization by Electrostatic Repulsions in the Two-Stranded α-Helical Coiled-Coil/Leucine Zipper. Protein Science 1995, 4 (2), 237–250. https://doi.org/10.1002/pro.5560040210.

(75) Baker, E. G.; Bartlett, G. J.; Crump, M. P.; Sessions, R. B.; Linden, N.; Faul, C. F. J.; Woolfson, D. N. Local and Macroscopic Electrostatic Interactions in Single α-Helices. Nature Chemical Biology 2015, 11 (3), 221–228. https://doi.org/10.1038/nchembio.1739.

(76) Huang, P.-S.; Oberdorfer, G.; Xu, C.; Pei, X. Y.; Nannenga, B. L.; Rogers, J. M.; DiMaio, F.; Gonen, T.; Luisi, B.; Baker, D. High Thermodynamic Stability of Parametrically Designed Helical Bundles. Science 2014, 346 (6208), 481–485. https://doi.org/10.1126/science.1257481.

(77) Egelman, E. H.; Xu, C.; DiMaio, F.; Magnotti, E.; Modlin, C.; Yu, X.; Wright, E.; Baker, D.; Conticello, V. P. Structural Plasticity of Helical Nanotubes Based on Coiled-Coil Assemblies. Structure 2015, 23 (2), 280–289. https://doi.org/10.1016/j.str.2014.12.008.

(78) Woolfson, D. N.; Alber, T. Predicting Oligomerization States of Coiled Coils. Protein Science 1995, 4 (8), 1596–1607. https://doi.org/10.1002/pro.5560040818.

(79) Woolfson, D. N.; Bartlett, G. J.; Burton, A. J.; Heal, J. W.; Niitsu, A.; Thomson, A. R.; Wood, C. W. De Novo Protein Design: How Do We Expand into the Universe of Possible Protein Structures? Current Opinion in Structural Biology 2015, 33, 16–26. https://doi.org/10.1016/j.sbi.2015.05.009.

(80) Niitsu Ai; Heal Jack W.; Fauland Kerstin; Thomson Andrew R.; Woolfson Derek N. Membrane-Spanning α-Helical Barrels as Tractable Protein-Design Targets. Philosophical Transactions of the Royal Society B: Biological Sciences 2017, 372 (1726), 20160213. https://doi.org/10.1098/rstb.2016.0213.

(81) Crick, F. H. The Fourier Transform of a Coiled-Coil. Acta crystallographica 1953, 6 (8–9), 685–689.

(82) Grigoryan, G.; DeGrado, W. F. Probing Designability via a Generalized Model of Helical Bundle Geometry. Journal of Molecular Biology 2011, 405 (4), 1079–1100. https://doi.org/10.1016/j.jmb.2010.08.058.

(83) Harbury, P. B.; Tidor, B.; Kim, P. S. Repacking Protein Cores with Backbone Freedom: Structure Prediction for Coiled Coils. PNAS 1995, 92 (18), 8408–8412. https://doi.org/10.1073/pnas.92.18.8408.

(84) Offer, G.; Sessions, R. Computer Modelling of the α-Helical Coiled Coil: Packing of Side-Chains in the Inner Core. Journal of Molecular Biology 1995, 249 (5), 967–987. https://doi.org/10.1006/jmbi.1995.0352.

(85) Wood, C. W.; Bruning, M.; Ibarra, A. Á.; Bartlett, G. J.; Thomson, A. R.; Sessions, R. B.; Brady, R. L.; Woolfson, D. N. CCBuilder: An Interactive Web-Based Tool for Building, Designing and Assessing Coiled-Coil Protein Assemblies. Bioinformatics 2014, 30 (21), 3029–3035. https://doi.org/10.1093/bioinformatics/btu502.

(86) Wood, C. W.; Woolfson, D. N. CCBuilder 2.0: Powerful and Accessible Coiled-Coil Modeling. Protein Science 2018, 27 (1), 103–111. https://doi.org/10.1002/pro.3279.

(87) Lupas, A. N.; Bassler, J. Coiled Coils – A Model System for the 21st Century. Trends in Biochemical Sciences 2017, 42 (2), 130–140. https://doi.org/10.1016/j.tibs.2016.10.007.

(88) Negron, C.; Keating, A. E. A Set of Computationally Designed Orthogonal Antiparallel Homodimers That Expands the Synthetic Coiled-Coil Toolkit. J. Am. Chem. Soc. 2014, 136 (47), 16544–16556. https://doi.org/10.1021/ja507847t.

(89) Jr, L. G.; Woolfson, D. N.; Alber, T. Buried Polar Residues and Structural Specificity in the GCN4 Leucine Zipper. Nature Structural & Molecular Biology 1996, 3 (12), 1011–1018. https://doi.org/10.1038/nsb1296-1011.

(90) Lumb, K. J.; Kim, P. S. A Buried Polar Interaction Imparts Structural Uniqueness in a Designed Heterodimeric Coiled Coil. Biochemistry 1998, 37 (37), 13042–13042. https://doi.org/10.1021/bi9850468.

(91) Fletcher, J. M.; Bartlett, G. J.; Boyle, A. L.; Danon, J. J.; Rush, L. E.; Lupas, A. N.; Woolfson, D. N. N@a and N@d: Oligomer and Partner Specification by Asparagine in Coiled-Coil Interfaces. ACS Chem. Biol. 2017, 12 (2), 528–538. https://doi.org/10.1021/acschembio.6b00935.

(92) Boyken, S. E.; Chen, Z.; Groves, B.; Langan, R. A.; Oberdorfer, G.; Ford, A.; Gilmore, J. M.; Xu, C.; DiMaio, F.; Pereira, J. H.; et al. De Novo Design of Protein Homo-Oligomers with Modular Hydrogen-Bond Network–Mediated Specificity. Science 2016, 352 (6286), 680–687. https://doi.org/10.1126/science.aad8865.

(93) Lizatović, R.; Aurelius, O.; Stenström, O.; Drakenberg, T.; Akke, M.; Logan, D. T.; André, I. A De Novo Designed Coiled-Coil Peptide with a Reversible PH-Induced Oligomerization Switch. Structure 2016, 24 (6), 946–955. https://doi.org/10.1016/j.str.2016.03.027.

(94) Ambroggio, X. I.; Kuhlman, B. Computational Design of a Single Amino Acid Sequence That Can Switch between Two Distinct Protein Folds. J. Am. Chem. Soc. 2006, 128 (4), 1154–1161. https://doi.org/10.1021/ja054718w.

(95) Aupič, J.; Lapenta, F.; Jerala, R. SwitCCh: Metal-Site Design for Controlling the Assembly of a Coiled-Coil Homodimer. ChemBioChem 2018, 19 (23), 2453–2457. https://doi.org/10.1002/cbic.201800578.

(96) Cerasoli, E.; Sharpe, B. K.; Woolfson, D. N. ZiCo:? A Peptide Designed to Switch Folded State upon Binding Zinc. J. Am. Chem. Soc. 2005, 127 (43), 15008–15009. https://doi.org/10.1021/ja0543604.

(97) Ciani, B.; Hutchinson, E. G.; Sessions, R. B.; Woolfson, D. N. A Designed System for Assessing How Sequence Affects α to β Conformational Transitions in Proteins. J. Biol. Chem. 2002, 277 (12), 10150–10155. https://doi.org/10.1074/jbc.M107663200.

(98) Slovic, A. m.; Lear, J. d.; DeGrado, W. f. De Novo Design of a Pentameric Coiled-Coil: Decoding the Motif for Tetramer versus Pentamer Formation in Water-Soluble Phospholamban*. The Journal of Peptide Research 2005, 65 (3), 312–321. https://doi.org/10.1111/j.1399-3011.2005.00244.x.

(99) Slovic, A. M.; Stayrook, S. E.; North, B.; DeGrado, W. F. X-Ray Structure of a Water-Soluble Analog of the Membrane Protein Phospholamban: Sequence Determinants Defining the Topology of Tetrameric and Pentameric Coiled Coils. Journal of Molecular Biology 2005, 348 (3), 777–787. https://doi.org/10.1016/j.jmb.2005.02.040.

(100) Spencer, R. K.; Hochbaum, A. I. The Phe-Ile Zipper: A Specific Interaction Motif Drives Antiparallel Coiled-Coil Hexamer Formation. Biochemistry 2017, 56 (40), 5300–5308. https://doi.org/10.1021/acs.biochem.7b00756.

(101) Spencer, R. K.; Hochbaum, A. I. X-Ray Crystallographic Structure and Solution Behavior of an Antiparallel Coiled-Coil Hexamer Formed by de Novo Peptides. Biochemistry 2016, 55 (23), 3214–3223. https://doi.org/10.1021/acs.biochem.6b00201.

(102) Ljubetič, A.; Lapenta, F.; Gradišar, H.; Drobnak, I.; Aupič, J.; Strmšek, Ž.; Lainšček, D.; Hafner-Bratkovič, I.; Majerle, A.; Krivec, N.; et al. Design of Coiled-Coil Protein-Origami Cages That Self-Assemble *in Vitro* and *in Vivo*. Nature Biotechnology 2017, 35 (11), 1094–1101. https://doi.org/10.1038/nbt.3994.

(103) Swainsbury, D. J. K.; Harniman, R. L.; Di Bartolo, N. D.; Liu, J.; Harper, W. F. M.; Corrie, A. S.; Jones, M. R. Directed Assembly of Defined Oligomeric Photosynthetic Reaction Centres through Adaptation with Programmable Extra-Membrane Coiled-Coil Interfaces. Biochim Biophys Acta 2016, 1857 (12), 1829–1839. https://doi.org/10.1016/j.bbabio.2016.09.002.

(104) Cristie-David, A. S.; Sciore, A.; Badieyan, S.; Escheweiler, J. D.; Koldewey, P.; Bardwell, J. C. A.; Ruotolo, B. T.; Marsh, E. N. G. Evaluation of de Novo-Designed Coiled Coils as off-the-Shelf Components for Protein Assembly. Mol. Syst. Des. Eng. 2017, 2 (2), 140–148. https://doi.org/10.1039/C7ME00012J.

(105) Sciore, A.; Su, M.; Koldewey, P.; Eschweiler, J. D.; Diffley, K. A.; Linhares, B. M.; Ruotolo, B. T.; Bardwell, J. C. A.; Skiniotis, G.; Marsh, E. N. G. Flexible, Symmetry-Directed Approach to Assembling Protein Cages. Proc Natl Acad Sci U S A 2016, 113 (31), 8681–8686. https://doi.org/10.1073/pnas.1606013113.

(106) Fletcher, J. M.; Harniman, R. L.; Barnes, F. R. H.; Boyle, A. L.; Collins, A.; Mantell, J.; Sharp, T. H.; Antognozzi, M.; Booth, P. J.; Linden, N.; et al. Self-Assembling Cages from Coiled-Coil Peptide Modules. Science 2013, 1226558. https://doi.org/10.1126/science.1233936.

