## Supplementary information for "Navigating the structural landscape of *de novo* α–helical bundles"

### Table of contents

SECTION 1 SEQUENCE LIST 3

SECTION 2 MATERIALS AND METHODS 4

SECTION 3 MATRIX-ASSISTED LASER DESORPTION/IONISATION - TIME OF FLIGHT (MALDI-TOF) AND ANALYTICAL HIGH-PRESSURE LIQUID CHROMATOGRAPHY (HPLC) 8

SECTION 4 CIRCULAR DICHROISM (CD) 12

SECTION 5 VARIABLE CONCENTRATION CIRCULAR DICHROISM 18

SECTION 6 ANALYTICAL ULTRACENTRIFUGATION 19

SECTION 7 CRYSTALLOGRAPHY TABLES 26

### Section 1 Sequence List

*Table S1 Sequences discussed in the article.*

| **Name** | **Sequence**<br>gabcdefgabcdefgabcdefgabcdef |
|---|---|
| CC-Hex | Ac-GELKAIAQELKAIAKELKAIAWELKAIAQGAG-$NH_2$ |
| CC-Hex-D24 | Ac-GELKAIAQELKAIAKELKAIAWEDKAIAQGAG-$NH_2$ |
| CC-Hex-L24D | Ac-GELKAIAQELKAIAKELKAIAWEDKAIAQG-$NH_2$ |
| CC-Hex-L24E | Ac-GELKAIAQELKAIAKELKAIAWEEKAIAQG-$NH_2$ |
| CC-Hex-L24K | Ac-GELKAIAQELKAIAKELKAIAWEKKAIAQG-$NH_2$ |
| CC-Hex-L24DAB | Ac-GELKAIAQELKAIAKELKAIAWEΓKAIAQG-$NH_2$ |
| CC-Hex-L24Nle | Ac-GELKAIAQELKAIAKELKAIAWEΔKAIAQG-$NH_2$ |
| CC-Hex-L24H | Ac-GELKAIAQELKAIAKELKAIAWEHKAIAQG-$NH_2$ |
| CC-Hex-LL | Ac-GELKALAQELKALAKELKALAWELKALAQG-$NH_2$ |
| CC-Hex-II | Ac-GEIKAIAQEIKAIAKEIKAIAWEIKAIAQG-$NH_2$ |
| CC-Hex-KgEb | Ac-GKLEAIAQKLEAIAKKLEAIAWKLEAIAQG-$NH_2$ |
| CC-Hex-KgEb_var | Ac-GKLEAIAQKLEAIAKKLEAIAWKLEAIAQGAG-$NH_2$ |
| CC-Hex-LL-KgEb | Ac-GKLEALAQKLEALAKKLEALAWKLEAIAQG-$NH_2$ |
| CC-AP-Tet | Ac-GELEALAQELEALAKKLKALAWKLKALAQG-$NH_2$ |

### Section 2 Materials and Methods

#### Peptide synthesis and purification

Peptides were synthesised by Fmoc methods on a CEM Liberty Blue automated solid-phase peptide synthesis apparatus with inline UV monitoring. Activation was achieved using DIC/Cl-HOBt. All peptides were produced as the C-terminal amide on a Rink amide ChemMatrix solid support or Rink Amide MBHA solid support, and N-terminally acetylated with 0.25 ml acetic anhydride and 0.3 ml pyridine in dimethylformamide (DMF). Cleavage from the support was effected with 25 ml trifluoroacetic acid (TFA) containing 0.4 ml triisopropylsilane and 0.4 ml water. The TFA solution was reduced to 5 ml under a flow of nitrogen. Crude peptides were precipitated with diethyl ether (45 ml) at 0 °C. The solid was recovered by centrifugation and redissolved in 1:1 acetonitrile:water before freeze-drying to yield crude peptides as white or pale yellow solids. Peptides were purified by reverse-phase HPLC with a gradient of acetonitrile in water (each containing 0.1% TFA) and, unless stated otherwise, over 30 min at room temperature. The stationary phase was a Phenomenex Luna 5 µm C18 column of dimensions 200 mm by 10 mm. Pure fractions were identified by analytical HPLC and MALDI mass spectrometry, and were pooled and freeze dried. Fmoc-protected proteinogenic amino acids, DMF and activators were purchased from AGTC Bioproduct. Fmoc-protected L-norleucine and L-2,4-diaminobutyric acid were purchased from Sigma Aldrich. All other solvents, Rink Amide ChemMatrix and Rink Amide MBHA solid support resin were purchased from Fisher Scientific, UK; PCAS Biomatrix, Canada; and Carbosynth Ltd, UK; respectively.

#### Analytical HPLC for designed sequences

Analytical HPLC was performed on Jasco 2000 series HPLC systems using a Phenomenex ''Kinetex'' 5 µm particle size, 100 Å pore size, C18 column of dimensions 100 × 4.6 mm. Chromatograms were monitored at 220 and 280 nm. Gradients were 20 to 80% or 40 to 100% acetonitrile in water (each containing 0.1% TFA) over 25 min.

#### MALDI-TOF mass spectrometry

MALDI-TOF mass spectra were collected on a Bruker UltraFlex MALDI-TOF mass spectrometer operating in positive-ion reflector mode. Peptides were spotted on a ground-steel target plate using dihydroxybenzoic acid as the matrix. Masses quoted are for the monoisotopic mass as the singly protonated species. Masses were measured to 0.1% accuracy.

#### Circular dichroism spectroscopy

Circular dichroism (CD) data were collected on a JASCO J-810 or J-815 spectropolarimeter fitted with a Peltier temperature controller. Unless stated otherwise, peptide samples were 10 or 50 µM solutions in phosphate-buffered saline (PBS, 8.2 mM sodium phosphate, 1.8 mM potassium phosphate, 137 mM sodium chloride, 2.7 mM potassium chloride at pH 7.4). pH titration experiments were conducted at 150 µM peptide concentration in 137 mM NaCl and 2.7 mM potassium chloride with the following buffer systems: pH 3–7, 50–100 mM citric acid/$Na_2HPO_4$ buffer. CD spectra were recorded in 5 or 1 mm path length quartz cuvettes at 5 °C. CD spectra were recorded with a scan rate of 100 nm $min^{-1}$, a 1 nm interval, a 1 nm bandwidth and a 1 s response time; and were an average of 8 scans recorded for the same sample. Thermal denaturation curves were acquired at 222 nm between 5 and 95 °C, with settings as above, a ramping rate of 40 °C per hour and are single recordings. Baselines recorded using the same buffer, cuvette and parameters were subtracted from each dataset. The spectra were converted from ellipticities (deg) to molar ellipticities (MRE, ($deg.cm^2.dmol^{-1}.res^{-1}$)) by normalising for concentration of peptide bonds and the cell path length. The N-terminal acetyl bond was included as a residue contributing to MRE but not the C-terminal amide.

#### Analytical ultracentrifugation

Analytical ultracentrifugation (AUC) was performed at 20 °C in a Beckman Optima XL-A or Beckman Optima XL-I analytical ultracentrifuge using an An-50 or An-60 Ti rotor. Unless stated otherwise, for sedimentation velocity experiments conducted at pH 7.4, solutions of 310 µl volume were in PBS at 150 µM peptide concentration, and placed in a sedimentation velocity cell with an epon two-channel centrepiece and quartz windows. The reference channel was loaded with 325 µl of buffer. The samples were centrifuged at 60 krpm, with absorbance scans taken across a radial range of 5.8 to 7.3 cm at 5 min intervals to a total of 120 scans. All data from a single run were fitted to a continuous c(s) distribution model using Sedfit at 95% confidence level or c(s,f/f0) distribution model using 95% confidence level with the amount of regularization in the s-direction, relative to the f/f0-direction, set to 1.0.[1] The partial specific volume ($\bar{v}$) for each of the peptides and the buffer densities and viscosities were calculated using Ultrascan II (http:/www.ultrascan.uthscsa.edu). Unless stated otherwise, solutions for sedimentation equilibrium experiments conducted at pH 7.4, were in PBS at 70 µM peptide concentration and to 110 µl per channel. Experiments were recorded in triplicate with a 6-channel centerpiece. Rotor speeds were in the range 20–48 krpm. Data were fitted to single, ideal species models using Ultrascan II. In all cases, 95% confidence limits were obtained by Monte Carlo analysis of the fits.

#### X-ray crystal structure determination

Freeze-dryed peptides were resuspended in deionised water to approximate concentrations of 10 mg $ml^{-1}$ for vapour-diffusion crystallisation trials using standard commercial screens (JCSG-plus™, Structure Screen 1 + 2, ProPlex™ and PACT Premier™) at 19 °C with 0.3 µl of the peptide solution equilibrated with 0.3 µl of the screen solution. Final crystallisation conditions for all peptides are provided in Table S2. To aid with cryoprotection, crystals were soaked in their respective reservoir solutions containing 25% glycerol prior to freezing. X-ray diffraction data were collected at the Diamond Light Source (Didcot, UK) on beamlines I02, I03, I04, I04-1 and I24, with the majority collected at wavelengths of 0.92 or 0.98 Å. Data were processed using the automated pipelines: Xia2 pipelines,[2] which ports data through DIALS[3] or MOSFLM[4] to POINTLESS and AIMLESS[5] as implemented in the CCP4 suite[6], or XDS to XSCALE[7]; or the AutoPROC pipelines[8], which use the same integrating and data reduction software in addition to STARANISO[9]. All structures were solved by molecular replacement using full or partial poly-alanine models (as dictated by the Matthews Coefficient), generated from existing coiled-coil structures, using PHASER[10]. Final structures were obtained after iterative rounds of model building with COOT[11] and refinement with PHENIX Refine[12] or REFMAC 5[13]. Late-stage models of all structures were submitted to PDB_REDO[14] and further refined with REFMAC 5. Solvent-exposed atoms lacking map density were either deleted or left at full occupancy. Data collection and refinement statistics are provided in Table S3.

#### Analogous structure search

A single biological assembly was generated for each crystal structure and submitted to the PDBeFold server (http://www.ebi.ac.uk/msd-srv/ssm/) searching the whole PDB archive as of 19/11/2018 without matching chain connectivity or matching to individual chains; and on 19/11/2018 the CAME TopSearch Server (https://topsearch.services.came.sbg.ac.at/) searching on the 18/4/2018 release of the PDB[67].

#### Bibliography


(1) Schuck, P. Size-Distribution Analysis of Macromolecules by Sedimentation Velocity Ultracentrifugation and Lamm Equation Modeling. *Biophysical Journal* **2000**, *78* (3), 1606–1619. https://doi.org/10.1016/S0006-3495(00)76713-0.

(2) Winter, G. Xia2: An Expert System for Macromolecular Crystallography Data Reduction. *J Appl Cryst* **2010**, *43* (1), 186–190. https://doi.org/10.1107/S0021889809045701.

(3) Winter, G.; Waterman, D. G.; Parkhurst, J. M.; Brewster, A. S.; Gildea, R. J.; Gerstel, M.; Fuentes-Montero, L.; Vollmar, M.; Michels-Clark, T.; Young, I. D.; et al. DIALS: Implementation and Evaluation of a New Integration Package. *Acta Cryst D* **2018**, *74* (2), 85–97. https://doi.org/10.1107/S2059798317017235.

(4) Powell, H. R. The Rossmann Fourier Autoindexing Algorithm in MOSFLM. *Acta Cryst D* **1999**, *55* (10), 1690–1695. https://doi.org/10.1107/S0907444999009506.

(5) Evans, P. R.; Murshudov, G. N. How Good Are My Data and What Is the Resolution? *Acta Cryst D* **2013**, *69* (7), 1204–1214. https://doi.org/10.1107/S0907444913000061.

(6) Winn, M. D.; Ballard, C. C.; Cowtan, K. D.; Dodson, E. J.; Emsley, P.; Evans, P. R.; Keegan, R. M.; Krissinel, E. B.; Leslie, A. G. W.; McCoy, A.; et al. Overview of the CCP4 Suite and Current Developments. *Acta Cryst D* **2011**, *67* (4), 235–242. https://doi.org/10.1107/S0907444910045749.

(7) Kabsch, W. XDS. *Acta Cryst D* **2010**, *66* (2), 125–132. https://doi.org/10.1107/S0907444909047337.

(8) Vonrhein, C.; Flensburg, C.; Keller, P.; Sharff, A.; Smart, O.; Paciorek, W.; Womack, T.; Bricogne, G. Data Processing and Analysis with the AutoPROC Toolbox. *Acta Cryst D* **2011**, *67* (4), 293–302. https://doi.org/10.1107/S0907444911007773.

(9) Tickle, I. J.; Flensburg, C.; Keller, P.; Paciorek, W.; Sharff, A.; Vonrhein, C.; Bricogne, G. STARANISO. *STARANISO. Cambridge, United Kingdom: Global Phasing Ltd.*

(10) McCoy, A. J.; Grosse-Kunstleve, R. W.; Adams, P. D.; Winn, M. D.; Storoni, L. C.; Read, R. J. Phaser Crystallographic Software. *J Appl Cryst* **2007**, *40* (4), 658–674. https://doi.org/10.1107/S0021889807021206.

(11) Emsley, P.; Lohkamp, B.; Scott, W. G.; Cowtan, K. Features and Development of Coot. *Acta Cryst D* **2010**, *66* (4), 486–501. https://doi.org/10.1107/S0907444910007493.

(12) Afonine, P. V.; Grosse-Kunstleve, R. W.; Echols, N.; Headd, J. J.; Moriarty, N. W.; Mustyakimov, M.; Terwilliger, T. C.; Urzhumtsev, A.; Zwart, P. H.; Adams, P. D. Towards Automated Crystallographic Structure Refinement with Phenix.Refine. *Acta Cryst D* **2012**, *68* (4), 352–367. https://doi.org/10.1107/S0907444912001308.

(13) Murshudov, G. N.; Vagin, A. A.; Dodson, E. J. Refinement of Macromolecular Structures by the Maximum-Likelihood Method. *Acta Cryst D* **1997**, *53* (3), 240–255. https://doi.org/10.1107/S0907444996012255.

(14) Joosten, R. P.; Long, F.; Murshudov, G. N.; Perrakis, A. The PDB_REDO Server for Macromolecular Structure Model Optimization. *IUCrJ* **2014**, *1* (4), 213–220. https://doi.org/10.1107/S2052252514009324.

### Section 3 Matrix-assisted laser desorption/ionisation - time of flight (MALDI-TOF) and Analytical high-pressure liquid chromatography (HPLC)

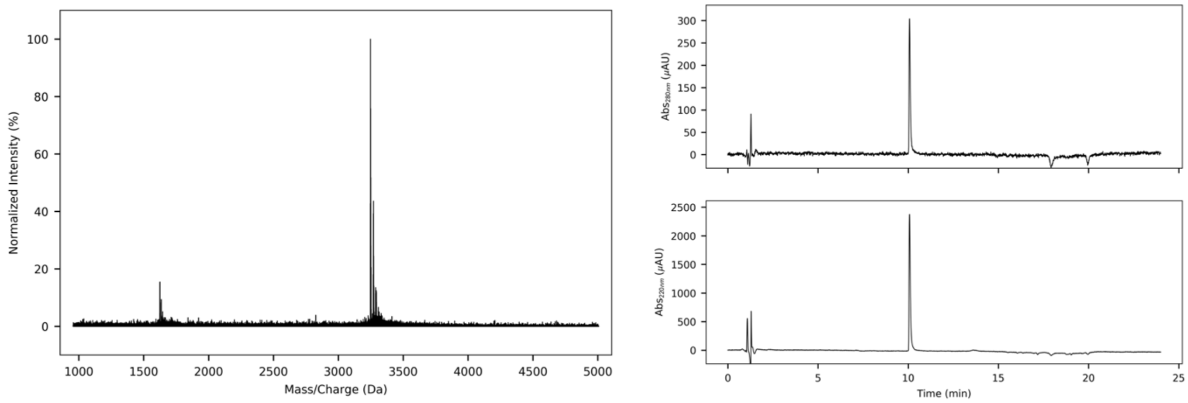


*Figure S3.1 CC-Hex-L24D MALDI-TOF MS (left) and HPLC traces from a gradient of 20 to 80% MeCN (0.1% TFA) in $H_2O$ (0.1% TFA) (right, 220 and 280 nm). Calculated average mass = 3248.8 Da, observed mass = 3248 Da [M+H$^+$].*

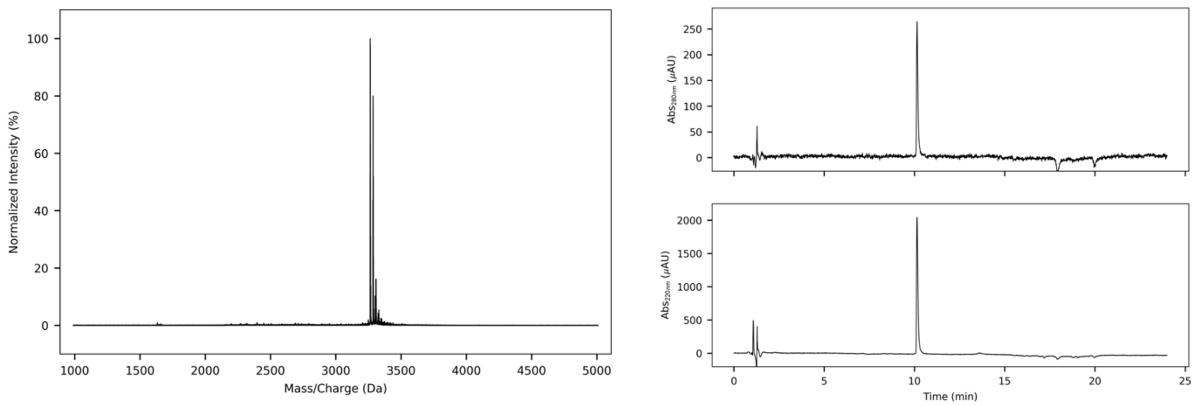


*Figure S3.2 CC-Hex-L24E MALDI-TOF MS (left) and HPLC traces from a gradient of 20 to 80% MeCN (0.1% TFA) in $H_2O$ (0.1% TFA) (right, 220 and 280 nm). Calculated average mass = 3262.8 Da, observed mass = 3264 Da [M+H$^+$].*

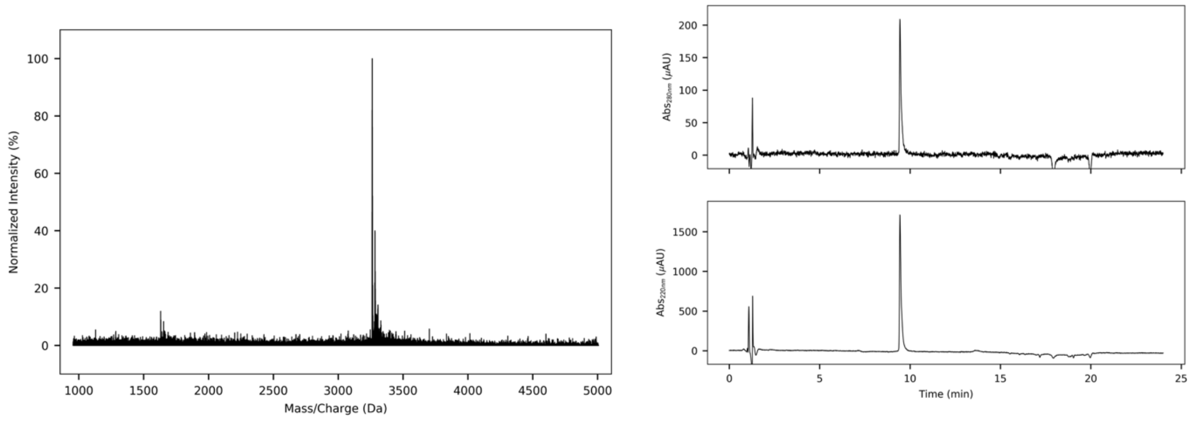


*Figure S3.3 CC-Hex-L24K MALDI-TOF MS (left) and HPLC traces from a gradient of 20 to 80% MeCN (0.1% TFA) in $H_2O$ (0.1% TFA) (right, 220 and 280 nm). Calculated average mass = 3261.9 Da, observed mass = 3263 Da [M+H$^+$].*

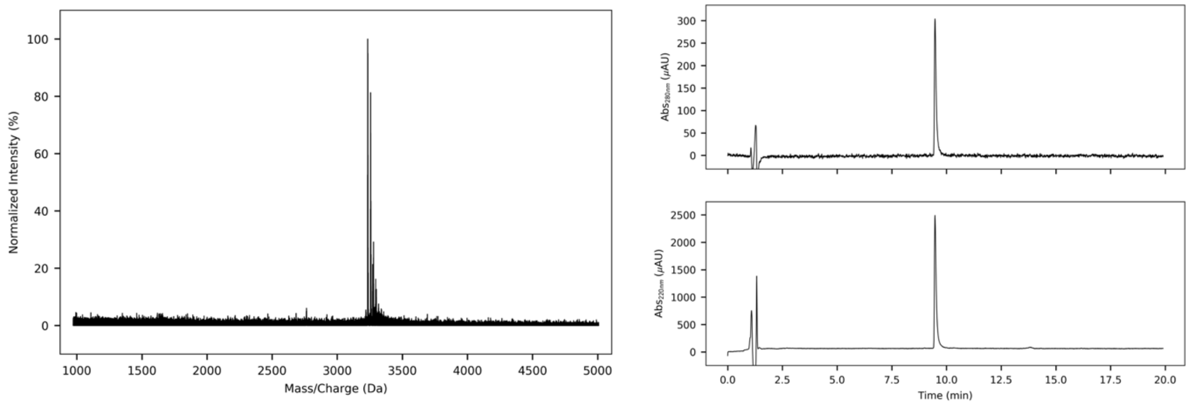


*Figure S3.4 CC-Hex-L24DAB MALDI-TOF MS (left) and HPLC traces from a gradient of 20 to 80% MeCN (0.1% TFA) in $H_2O$ (0.1% TFA) (right, 220 and 280 nm). Calculated average mass = 3233.9 Da, observed mass = 3234 Da [$M+H^+$].*

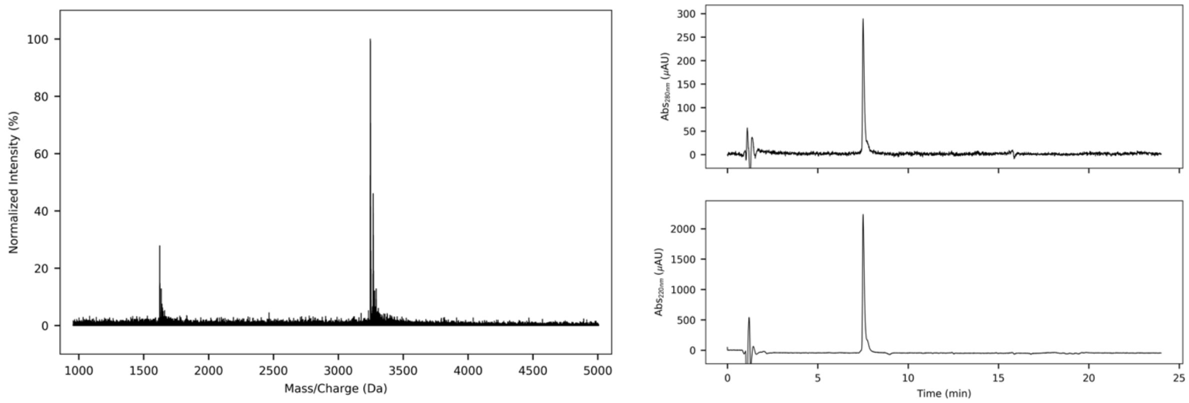


*Figure S3.5 CC-Hex-L24Nle MALDI-TOF MS (left) and HPLC traces from a gradient of 40 to 100% MeCN (0.1% TFA) in $H_2O$ (0.1% TFA) (right, 220 and 280 nm). Calculated average mass = 3246.8 Da, observed mass = 3247 Da [$M+H^+$].*

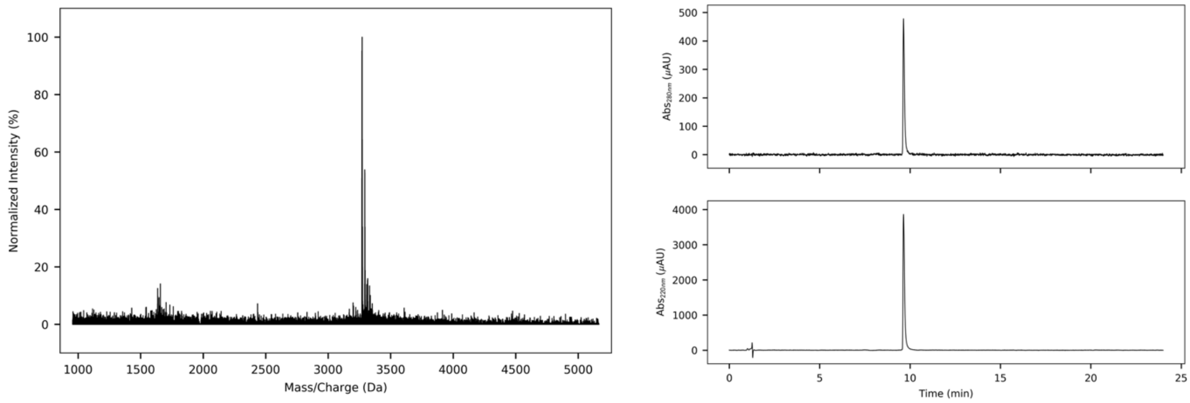


*Figure S3.6 CC-Hex-L24H MALDI-TOF MS (left) and HPLC traces from a gradient of 20 to 80% MeCN (0.1% TFA) in $H_2O$ (0.1% TFA) (right, 220 and 280 nm). Calculated average mass = 3270.8 Da, observed mass = 3272 Da [$M+H^+$].*

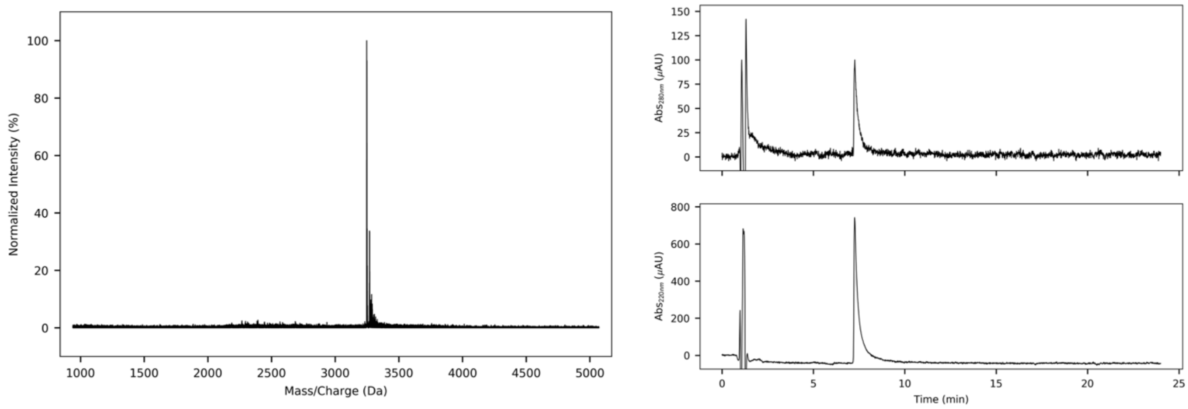


*Figure S3.7 CC-Hex-LL MALDI-TOF MS (left) and HPLC traces from a gradient of 40 to 100% MeCN (0.1% TFA) in $H_2O$ (0.1% TFA) (right, 220 and 280 nm). Calculated average mass = 3246.8 Da, observed mass = 3249 Da [$M+H^+$].*

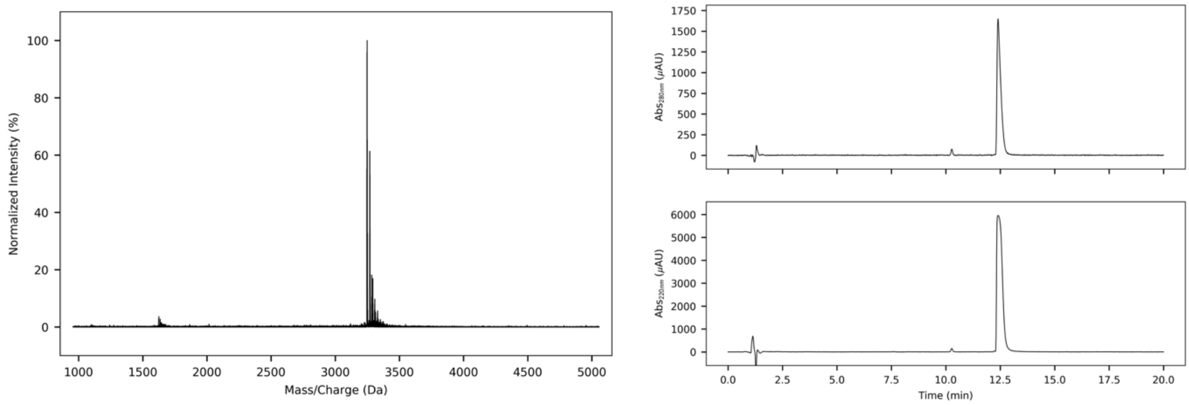


*Figure S3.8 CC-Hex-II MALDI-TOF MS (left) and HPLC traces from a gradient of 20 to 80% MeCN (0.1% TFA) in $H_2O$ (0.1% TFA) (right, 220 and 280 nm). Calculated average mass = 3246.8 Da, observed mass = 3248 Da [$M+H^+$].*

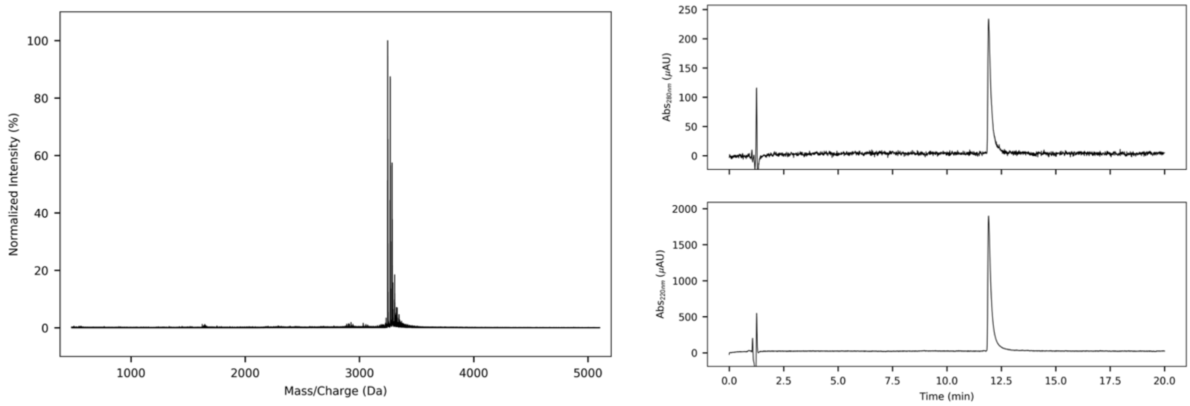


*Figure S3.9 CC-Hex-LL-KgEb MALDI-TOF MS (left) and HPLC traces from a gradient of 20 to 80% MeCN (0.1% TFA) in $H_2O$ (0.1% TFA) (right, 220 and 280 nm). Calculated average mass = 3246.8 Da, observed mass = 3247 Da [$M+H^+$].*

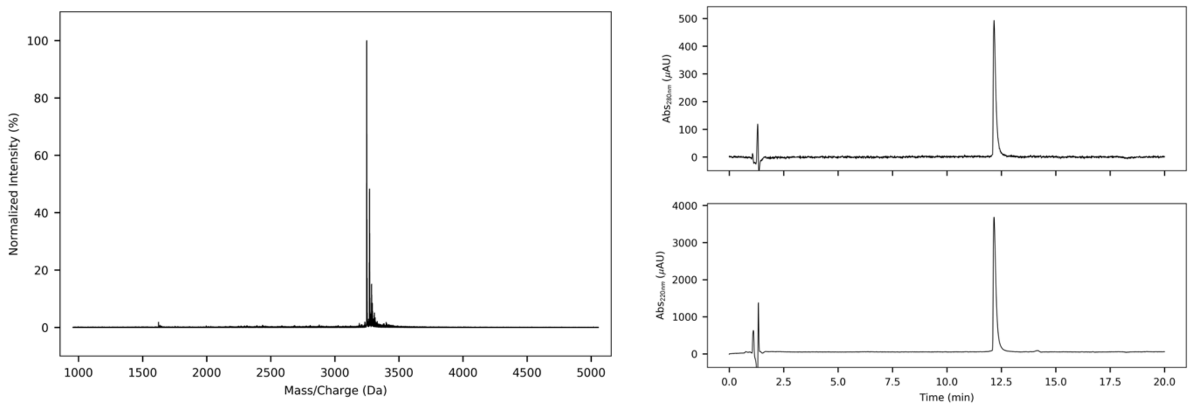


*Figure S3.10 CC-Hex-KgEb MALDI-TOF MS (left) and HPLC traces from a gradient of 20 to 80% MeCN (0.1% TFA) in $H_2O$ (0.1% TFA) (right, 220 and 280 nm). Calculated average mass = 3246.8 Da, observed mass = 3249 Da [$M+H^+$].*

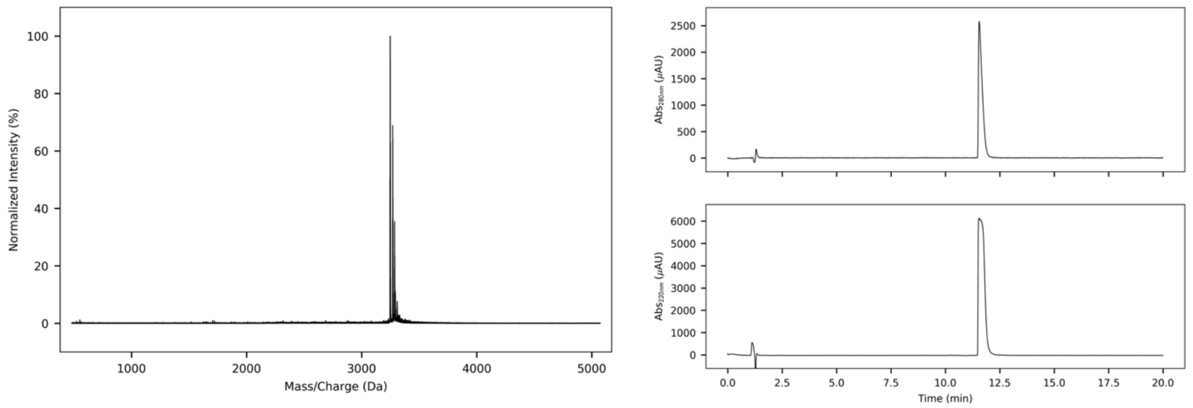


*Figure S3.11 apCC-Tet MALDI-TOF MS (left) and HPLC traces from a gradient of 20 to 80% MeCN (0.1% TFA) in $H_2O$ (0.1% TFA) (right, 220 and 280 nm). Calculated average mass = 3246.8 Da, observed mass = 3248 Da.*

### Section 4 Circular dichroism (CD)

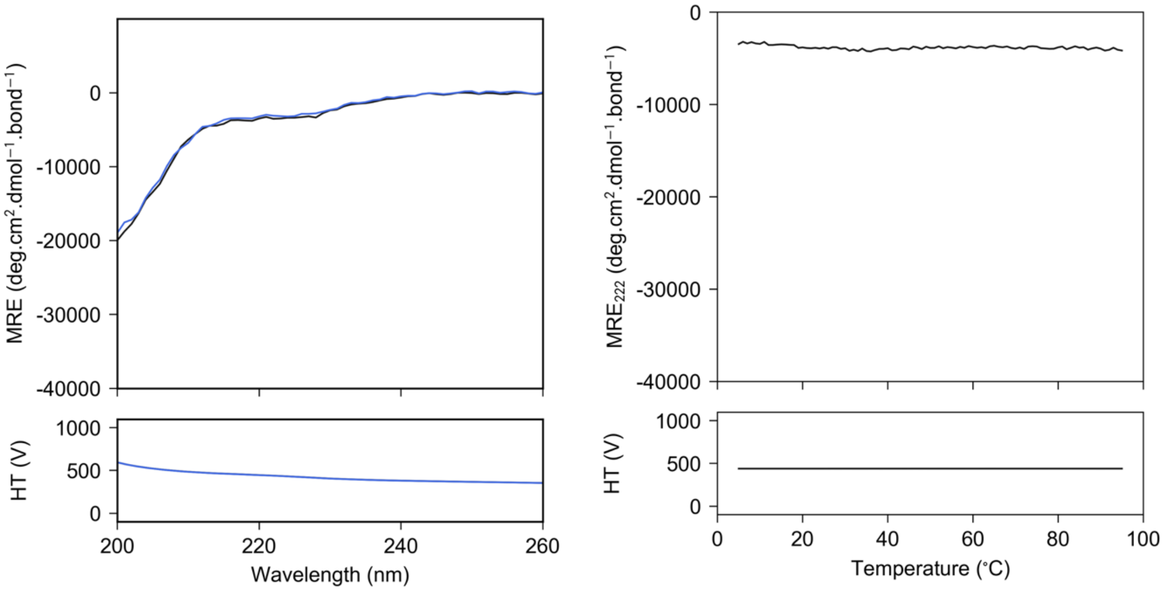


*Figure S4.1 CC-Hex-L24D CD spectrum at 5 °C before heating and after cooling (left top, black and blue respectively) and thermal denaturation profile monitored at 222 nm (top right). Plot of the high-tension voltage applied to the detector (left and right, bottom). Conditions: 50 μM peptide concentration, PBS (pH 7.4).*

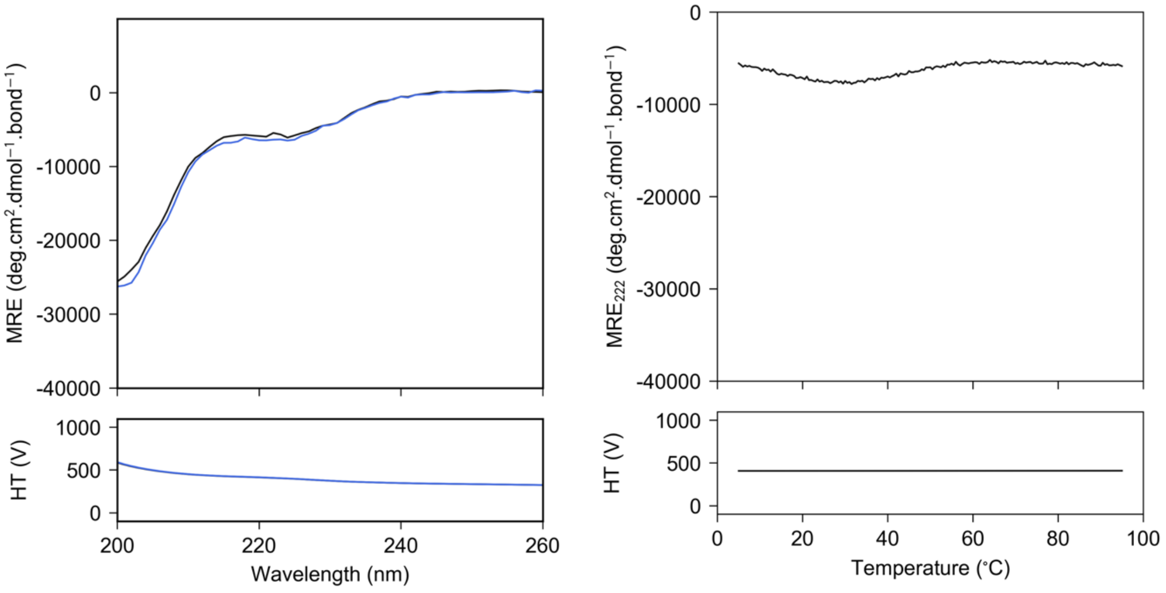


*Figure S4.2 CC-Hex-L24E CD spectrum at 5 °C before heating and after cooling (left top, black and blue respectively) and thermal denaturation profile monitored at 222 nm (top right). Plot of the high-tension voltage applied to the detector (left and right, bottom). Conditions: 50 μM peptide concentration, PBS (pH 7.4).*

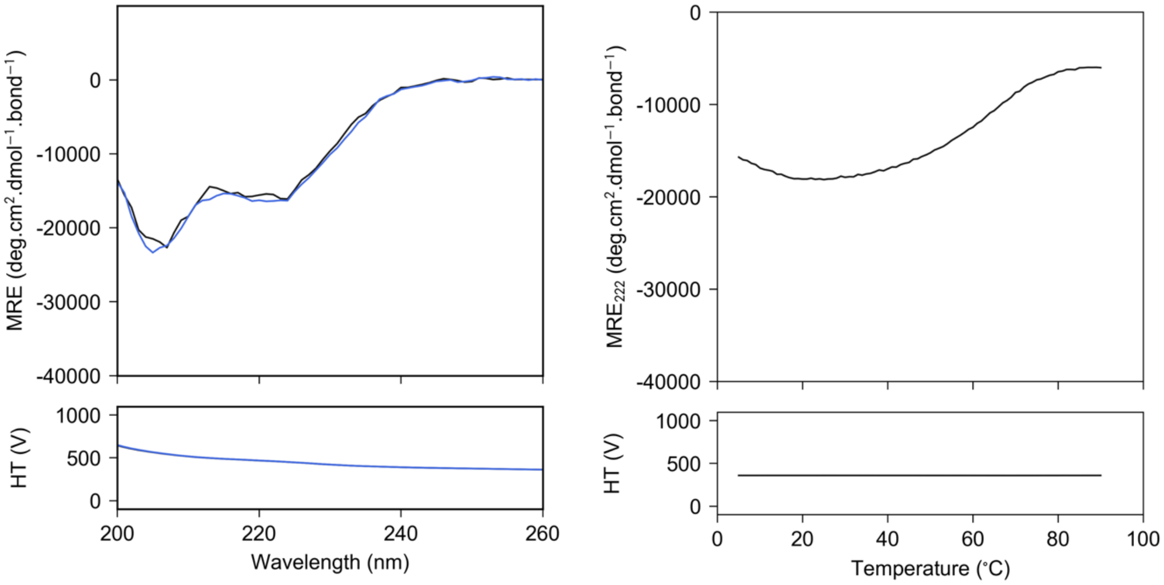


*Figure S4.3 CC-Hex-L24K CD spectrum at 5 °C before heating and after cooling (left top, black and blue respectively) and thermal denaturation profile monitored at 222 nm (top right). Plot of the high-tension voltage applied to the detector (left and right, bottom). Conditions: 50 μM peptide concentration, PBS (pH 7.4).*

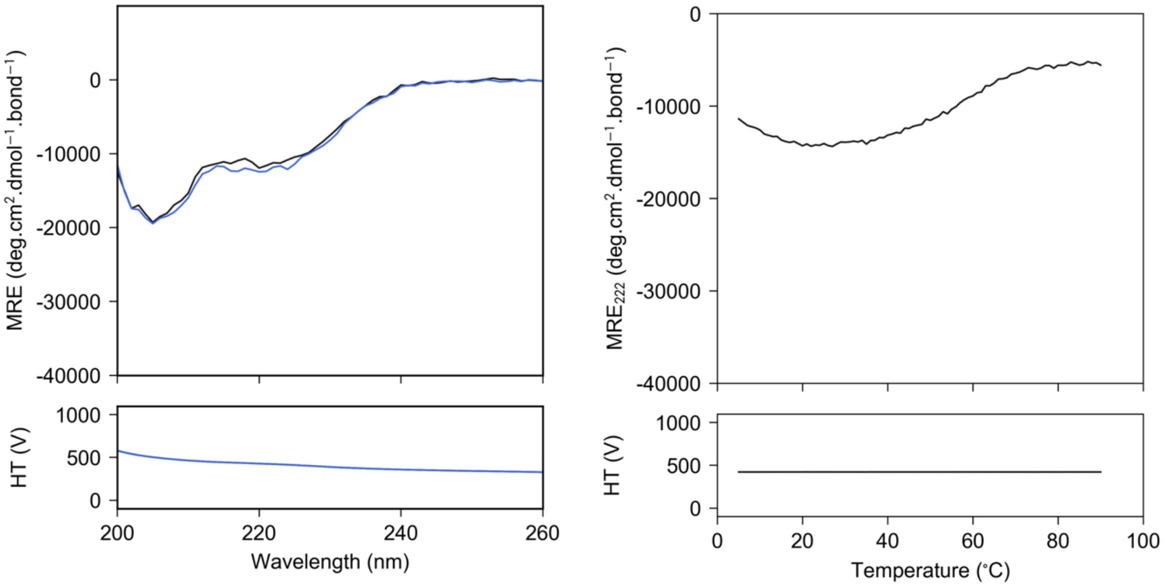


*Figure S4.4 CC-Hex-L24DAB CD spectrum at 5 °C before heating and after cooling (left top, black and blue respectively) and thermal denaturation profile monitored at 222 nm (top right). Plot of the high-tension voltage applied to the detector (left and right, bottom). Conditions: 50 μM peptide concentration, PBS (pH 7.4).*

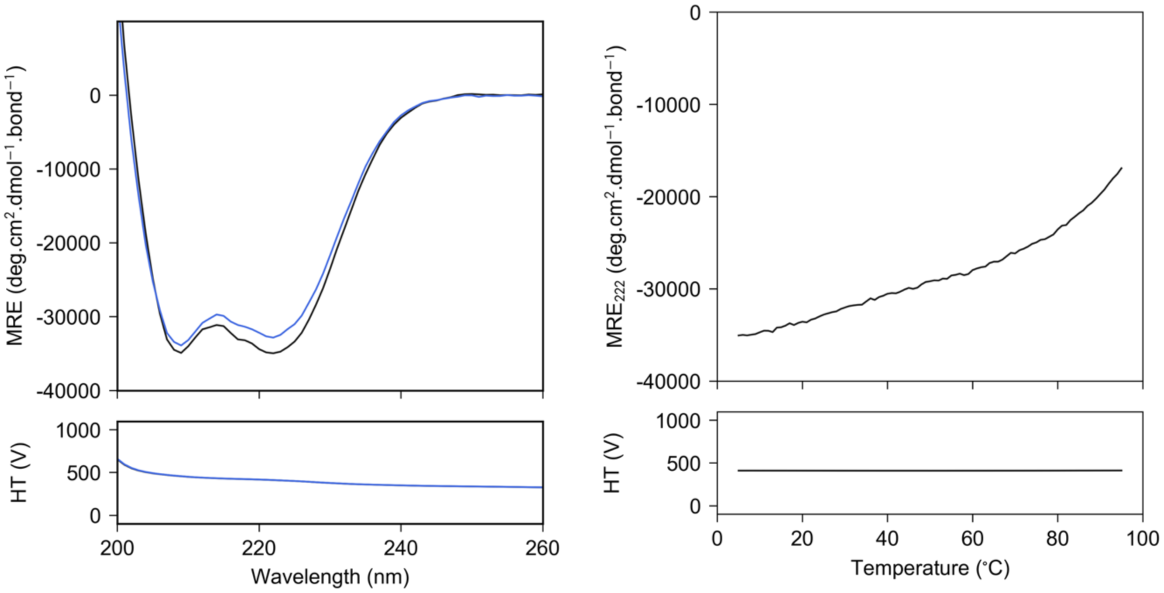


*Figure S4.5 CC-Hex-L24Nle CD spectrum at 5 °C before heating and after cooling (left top, black and blue respectively) and thermal denaturation profile monitored at 222 nm (top right). Plot of the high-tension voltage applied to the detector (left and right, bottom). Conditions: 10 μM peptide concentration, PBS (pH 7.4).*

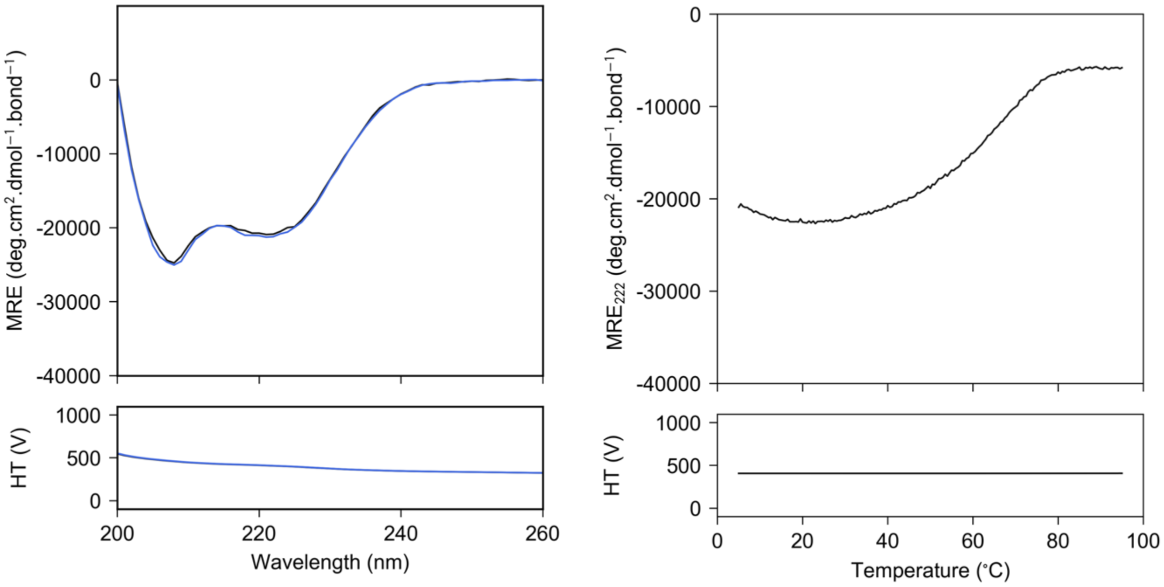


*Figure S4.6 CC-Hex-L24H CD spectrum at 5 °C before heating and after cooling (left top, black and blue respectively) and thermal denaturation profile monitored at 222 nm (top right). Plot of the high-tension voltage applied to the detector (left and right, bottom). Conditions: 50 μM peptide concentration, PBS (pH 7.4).*

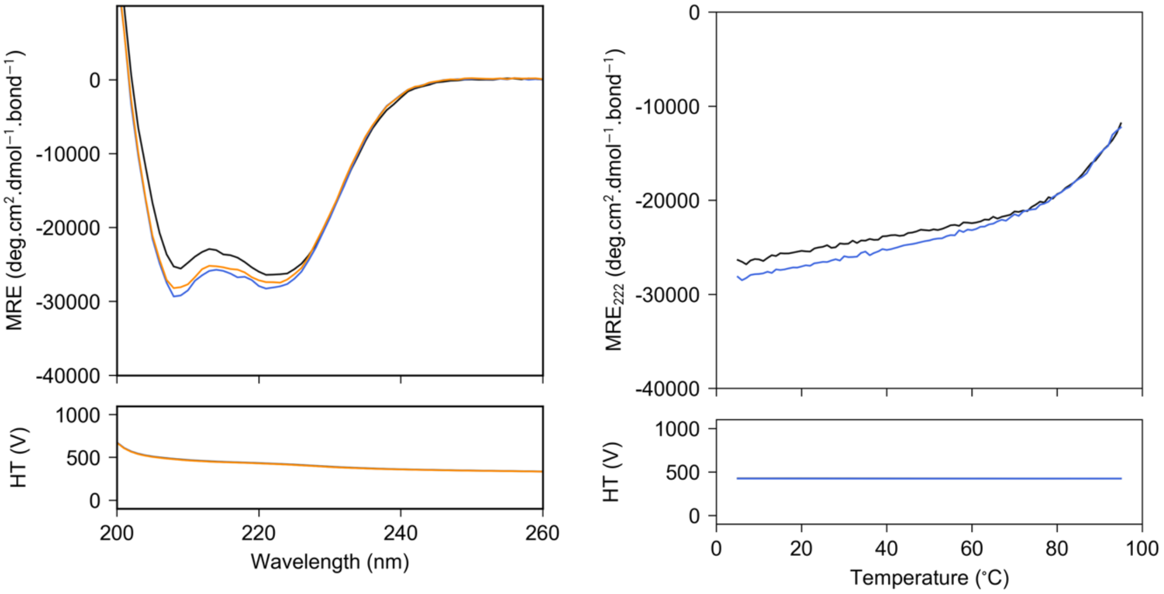


*Figure S4.7 CC-Hex-LL CD spectrum at 5 °C before heating, after cooling and after heating and cooling twice (left top, black, blue and orange respectively) and thermal denaturation profile and repeated thermal denaturation profile after cooling monitored at 222 nm (top right, black and blue respectively). Plot of the high-tension voltage applied to the detector (left and right, bottom). Conditions: 10 μM peptide concentration, PBS (pH 7.4).*

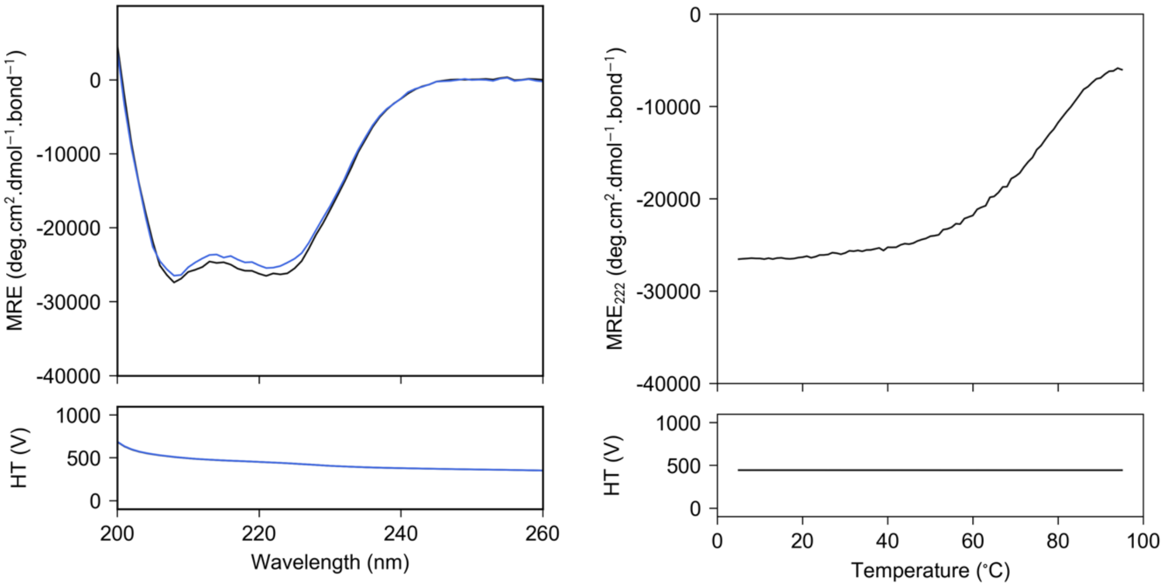


*Figure S4.8 CC-Hex-II CD spectrum at 5 °C before heating and after cooling (left top, black and blue respectively) and thermal denaturation profile monitored at 222 nm (top right). Plot of the high-tension voltage applied to the detector (left and right, bottom). Conditions: 10 μM peptide concentration, PBS (pH 7.4).*

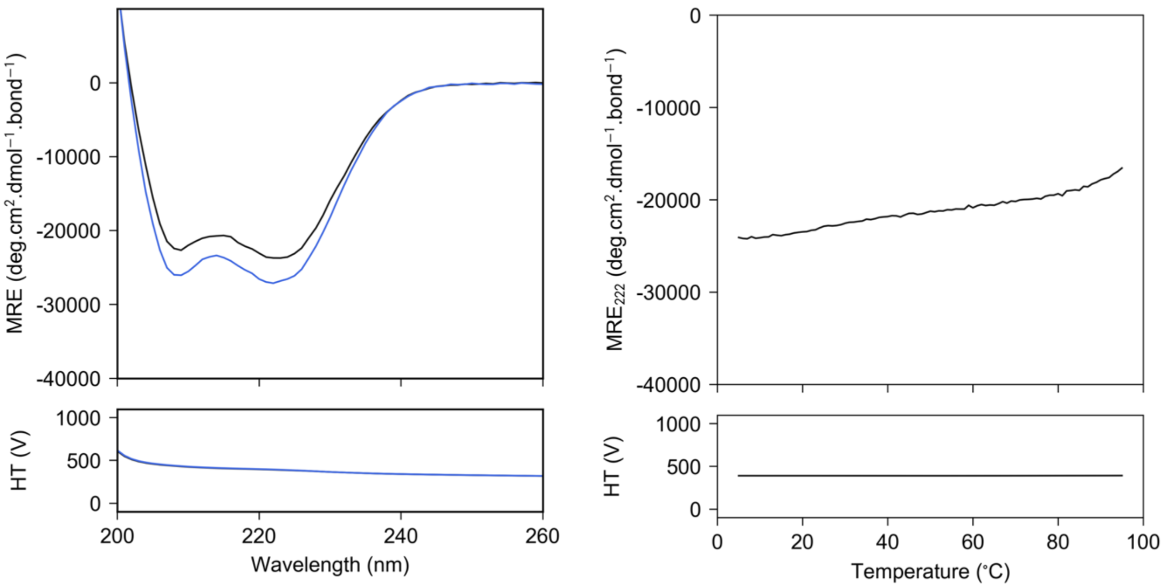


*Figure S4.9 CC-Hex-LL-KgEb CD spectrum at 5 °C before heating and after cooling (left top, black and blue respectively) and thermal denaturation profile monitored at 222 nm (top right). Plot of the high-tension voltage applied to the detector (left and right, bottom). Conditions: 10 μM peptide concentration, PBS (pH 7.4).*

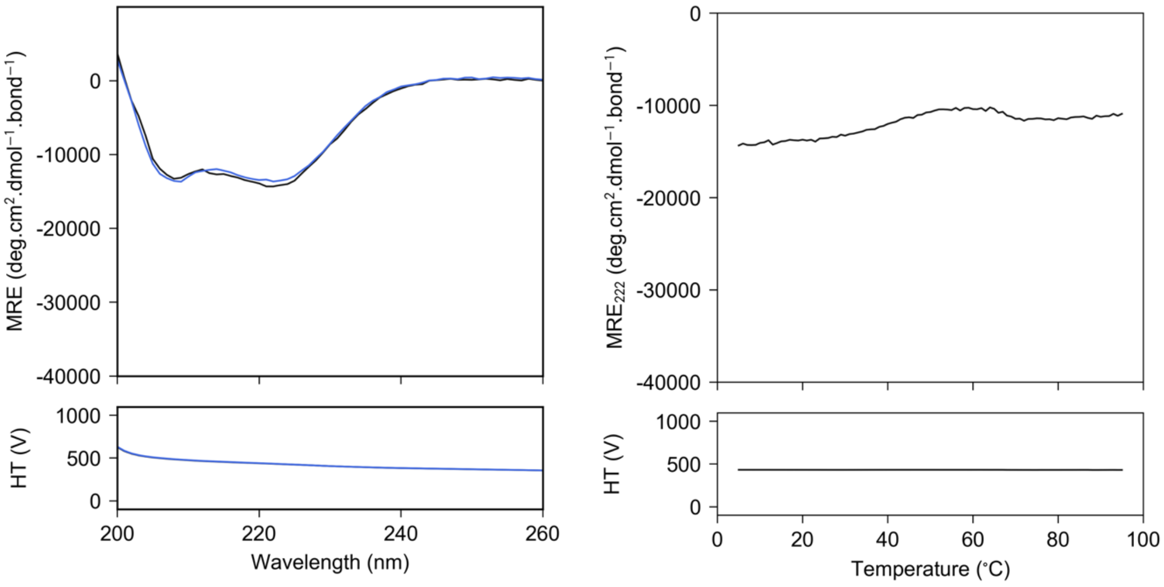


*Figure S4.10 CC-Hex-KgEb CD spectrum at 5 °C before heating and after cooling (left top, black and blue respectively) and thermal denaturation profile monitored at 222 nm (top right). Plot of the high-tension voltage applied to the detector (left and right, bottom). Conditions: 10 μM peptide concentration, PBS (pH 7.4).*

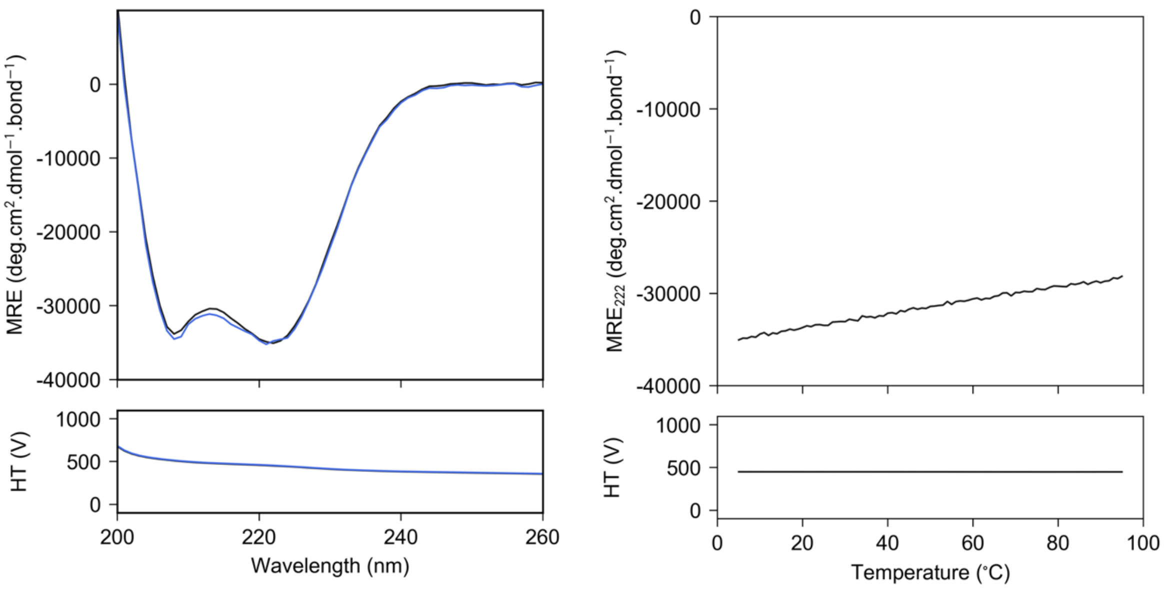


*Figure S4.11 apCC-Tet CD spectrum at 5 °C before heating and after cooling (left top, black and blue respectively) and thermal denaturation profile monitored at 222 nm (top right). Plot of the high-tension voltage applied to the detector (left and right, bottom). Conditions: 10 μM peptide concentration, PBS (pH 7.4).*

### Section 5 Variable concentration circular dichroism

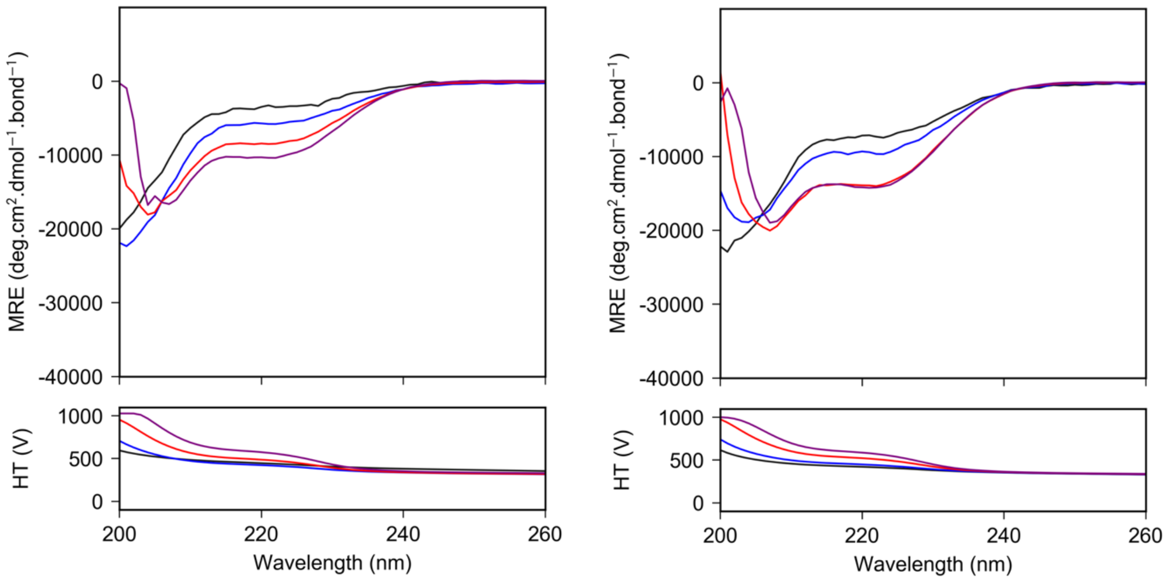


*Figure S5.1 Variable concentration CD spectrum at 20 °C for CC-Hex-L24D (left) and CC-Hex-L24E (right). Colours black, blue, red and purple indicate 50 $\mu$M, 100 $\mu$M, 200 $\mu$M and 300 $\mu$M peptide concentration respectively. Plot of the high-tension voltage applied to the detector (left and right, bottom). Conditions: PBS (pH 7.4).*

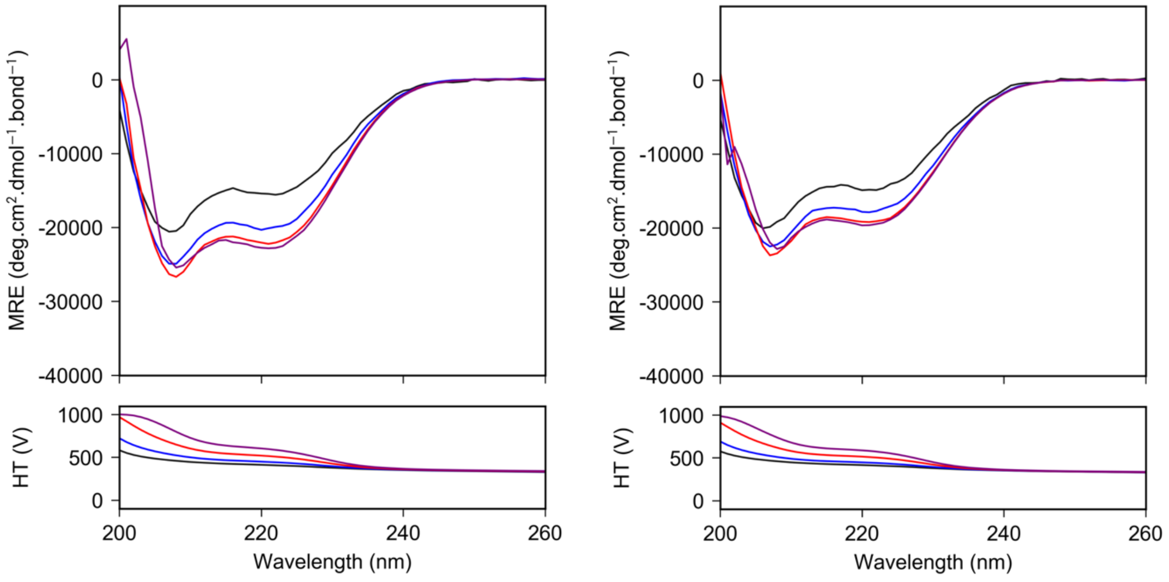


*Figure S5.2 Variable concentration CD spectrum at 20 °C for CC-Hex-L24K (left) and CC-Hex-L24DAB (right). Colours black, blue, red and purple indicate 50 $\mu$M, 100 $\mu$M, 200 $\mu$M and 300 $\mu$M peptide concentration respectively. Plot of the high-tension voltage applied to the detector (left and right, bottom). Conditions: PBS (pH 7.4).*

### Section 6 Analytical Ultracentrifugation

No data was collected for the sequence CC-Hex-KgEb as it aggregates at low concentration (10 μM) when span to 3,000 rpm.

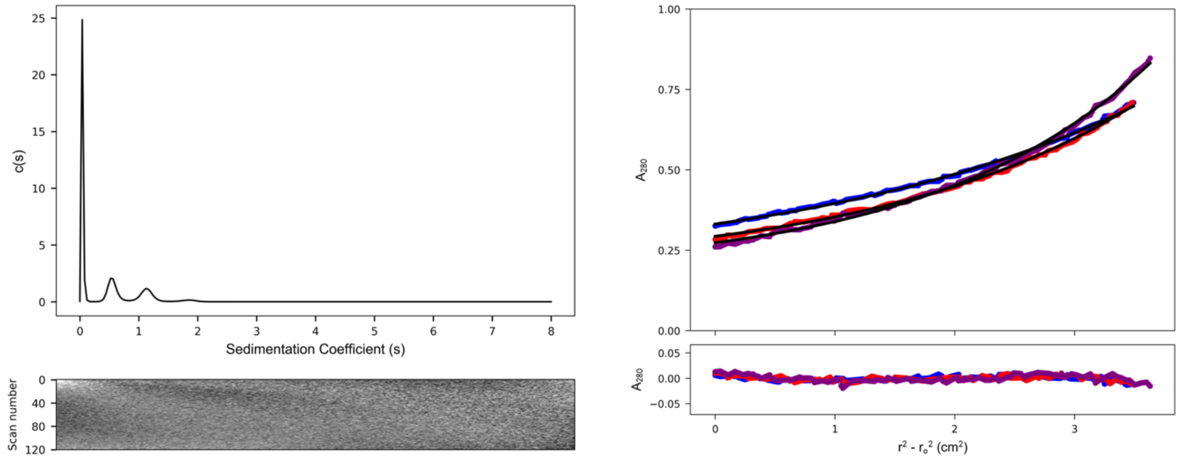


*Figure S6.1 CC-Hex-L24D AUC data and fits (top) and residuals (bottom) ($\bar{v}$ = 0.766 $cm^3$ $g^{-1}$). Left: continuous c(s) distribution from sedimentation-velocity data at 60k rpm returning two main peaks with an f/f0 = 1.586: first peak has s = 0.559 S, s20,w = 0.668 S and Mw = 5,157 Da (1.6 × monomer mass); and second peak has s = 1.135 S, s20,w = 1.356 S and Mw = 14,927 Da (4.6 × monomer mass) at 95% confidence limit. Conditions: 150 μM peptide concentration, PBS (pH 7.4). Right: sedimentation-equilibrium data (top, dots) and fitted single-ideal species model curves at 29k (blue), 33k (red) and 36k (purple) rpm. The fit returns a mass of 6,702 Da (2.1 × monomer mass, 95% confidence limits 6,617 – 6,787). Bottom: residuals for the, above fits using the same colour scheme as above. Conditions: 70 μM peptide concentration, PBS (pH 7.4).*

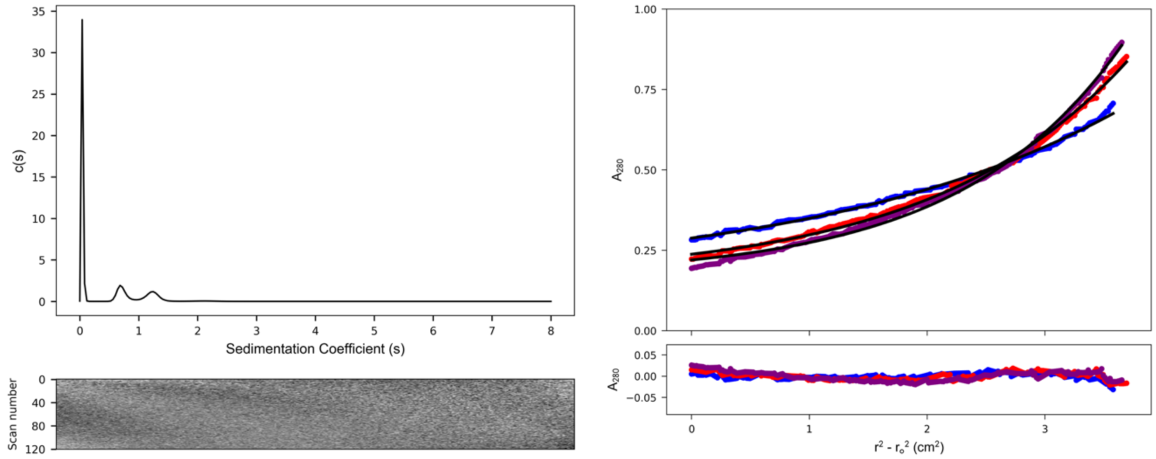


*Figure S6.2 CC-Hex-L24E AUC data and fits (top) and residuals (bottom) ($\bar{v}$ = 0.768 $cm^3$ $g^{-1}$). Left: continuous c(s) distribution from sedimentation-velocity data at 60k rpm returning two main peaks with an f/f0 = 1.433: first peak has s = 0.722 S, s20,w = 0.868 S and Mw = 6,580 Da (2.0 × monomer mass); and second peak has s = 1.233 S, s20,w = 1.484 S and Mw = 14,696 Da (4.5 × monomer mass) at 95% confidence limit. Conditions: 150 μM peptide concentration, PBS (pH 7.4). Right: sedimentation-equilibrium data (top, dots) and fitted single-ideal species model curves at 32k (blue), 40k (red), and 44k (purple) rpm. The fit returns a mass of 7,314 Da (2.2 × monomer mass, 95% confidence limits 7,252 – 7,375). Bottom: residuals for the, above fits using the same colour scheme as above. Conditions: 70 μM peptide concentration, PBS (pH 7.4).*

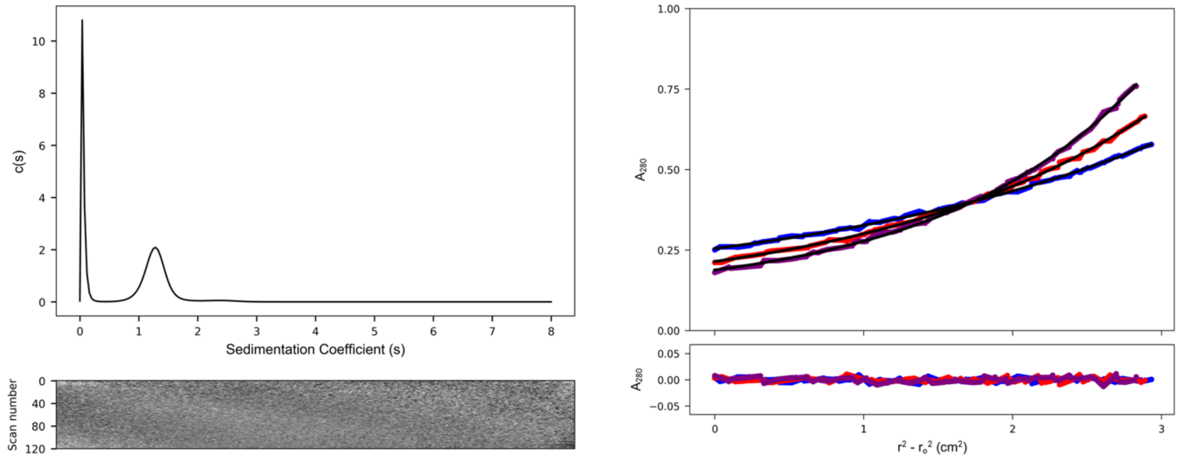


*Figure S6.3 CC-Hex-L24K AUC data and fits (top) and residuals (bottom) ($\bar{v}$ = 0.775 $cm^3$ $g^{-1}$). Left: continuous c(s) distribution from sedimentation-velocity data at 60k rpm returning s = 1.279 S, s20,w = 1.596 S, f/f0 = 1.297 and mw = 14,168 Da (4.3 × monomer mass) at 95% confidence level. Conditions: 150 µM peptide concentration, PBS (pH 7.4). Right: sedimentation-equilibrium data (top, dots) and fitted single-ideal species model curves at 28k (blue), 36k (red) and 44k (purple) rpm. The fit returns a mass of 15,320 Da (4.7 × monomer mass, 95% confidence limits 15,235 – 15,400). Bottom: residuals for the above fits using the same colour scheme as above. Conditions: 70 µM peptide concentration, PBS (pH 7.4).*

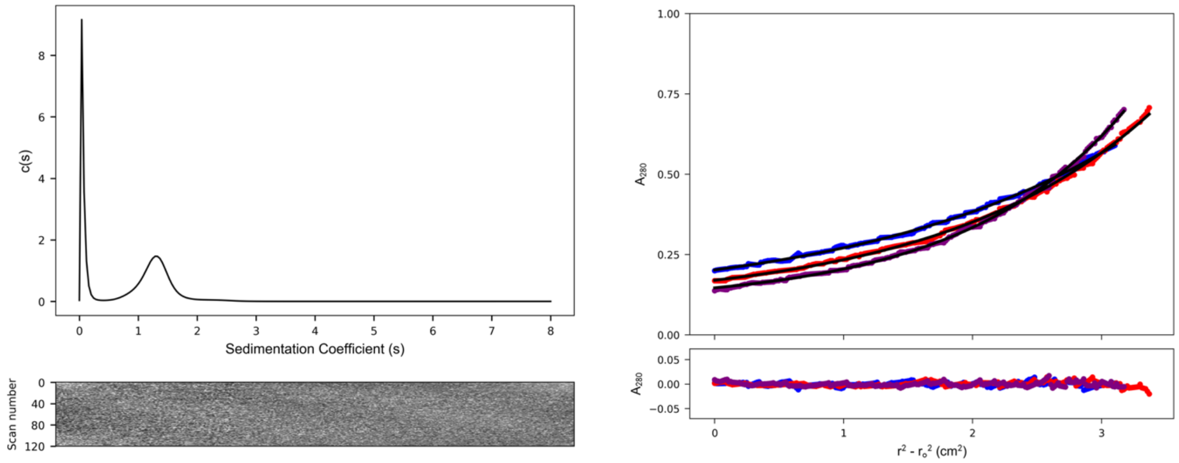


*Figure S6.4 CC-Hex-L24DAB AUC data and fits (top) and residuals (bottom) ($\bar{v}$ = 0.746 $cm^3$ $g^{-1}$). Left: continuous c(s) distribution from sedimentation-velocity data at 60k rpm returning s = 1.305 S, s20,w = 1.616 S, f/f0 = 1.334 and mw = 15,065 Da (4.7 × monomer mass) at 95% confidence level. Conditions: 150 µM peptide concentration, PBS (pH 7.4). Right: sedimentation-equilibrium data (top, dots) and fitted single-ideal species model curves at 26k (blue), 29k (red) and 33k (purple) rpm. The fit returns a mass of 14,580 Da (4.5 × monomer mass, 95% confidence limits 14,508 – 14,645). Bottom: residuals for the above fits using the same colour scheme as above. Conditions: 70 µM peptide concentration, PBS (pH 7.4).*

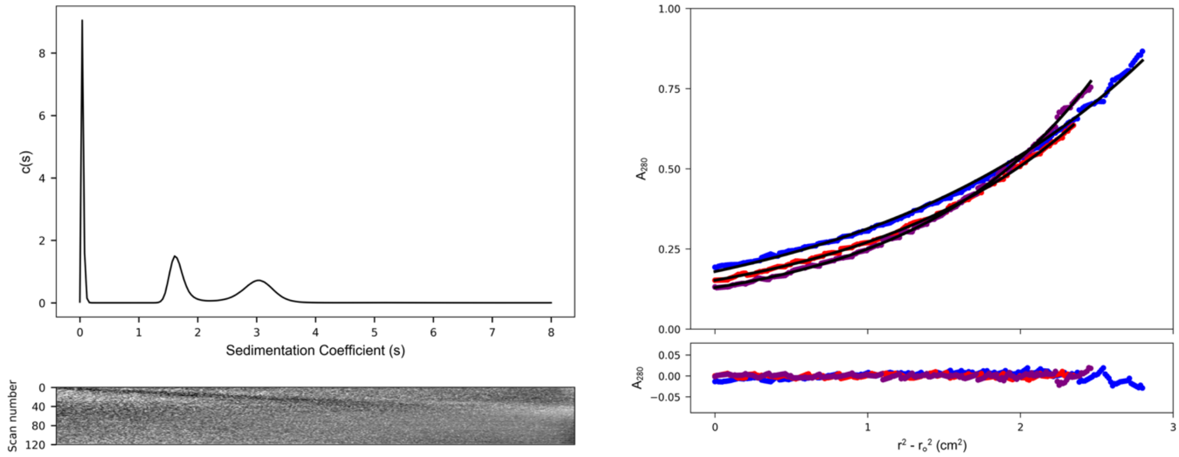


*Figure S6.5 CC-Hex-L24Nle AUC data and fits (top) and residuals (bottom) ($\bar{v}$ = 0.772 cm$^3$ g$^{-1}$). Left: continuous c(s) distribution from sedimentation-velocity data at 60k rpm returning two main peaks with an f/f0 = 1.108: first peak has s = 1.673 S, s20,w = 2.046 S and Mw = 16,203 Da (5.0 × monomer mass); and second peak has s = 3.004 S, s20,w = 3.674 S and Mw = 38,988 Da (12 × monomer mass) at 95% confidence limit. Conditions: 150 µM peptide concentration, PBS (pH 7.4). Right: sedimentation-equilibrium data (top, dots) and fitted single-ideal species model curves at 23k (blue), 26k (red) and 29k (purple) rpm. The fit returns a mass of 20,160 Da (6.2 × monomer mass, 95% confidence limits 19,893 – 20,438). Bottom: residuals for the above fits using the same colour scheme as above. Conditions: 70 µM peptide concentration, PBS (pH 7.4).*

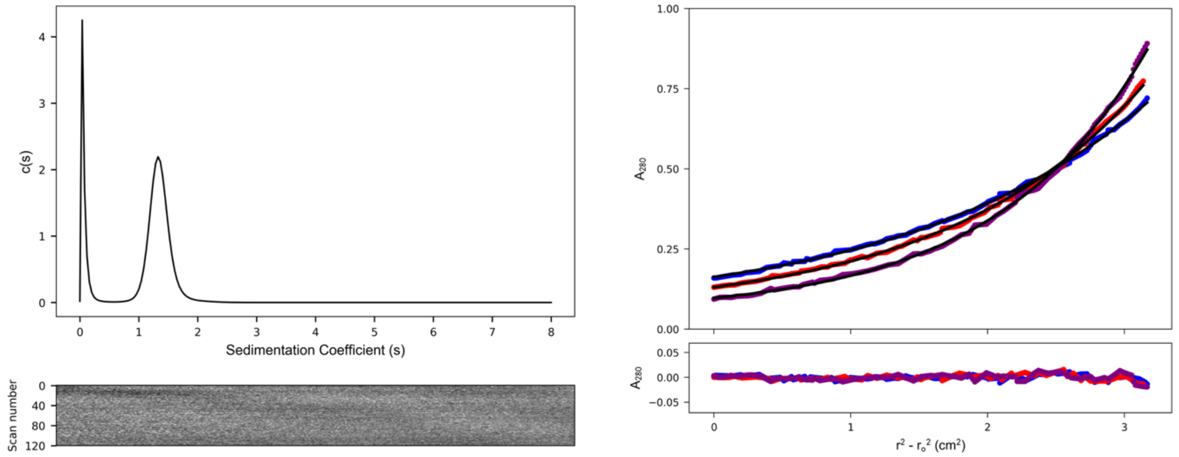


*Figure S6.6 CC-Hex-L24H AUC data and fits (top) and residuals (bottom) ($\bar{v}$ = 0.768 cm$^3$ g$^{-1}$). Left: continuous c(s) distribution from sedimentation-velocity data at 60k rpm returning s = 1.351 S, s20,w = 1.627 S, f/f0 = 1.244 and mw = 13,645 Da (4.2 × monomer mass) at 95% confidence level. Conditions: 150 µM peptide concentration, PBS (pH 7.4). Right: sedimentation-equilibrium data (top, dots) and fitted single-ideal species model curves at 26k (blue), 29k (red) and 33k (purple) rpm. The fit returns a mass of 16,040 Da (4.9 × monomer mass, 95% confidence limits 15,983 – 16,103). Bottom: residuals for the above fits using the same colour scheme as above. Conditions: 70 µM peptide concentration, PBS (pH 7.4).*

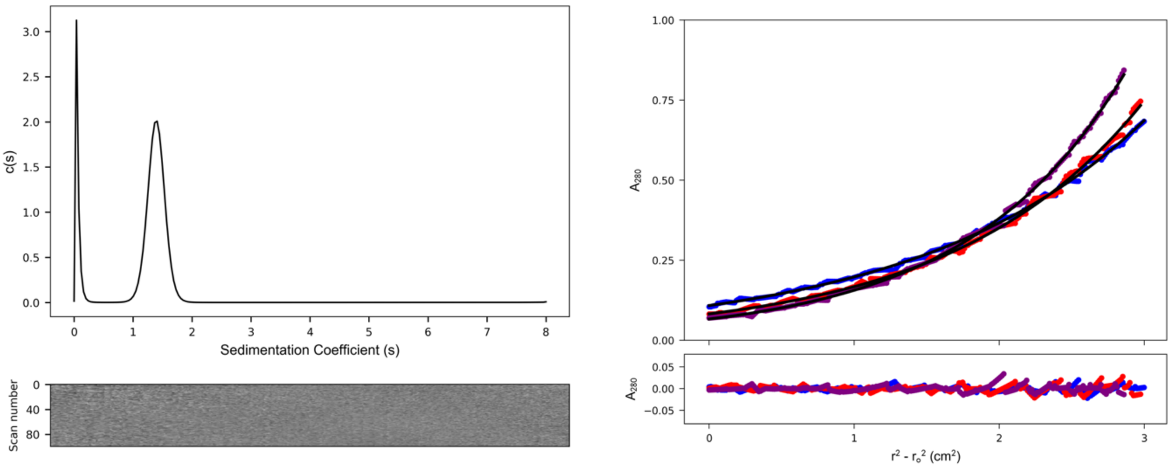


*Figure S6.7 CC-Hex-LL AUC data and fits (top) and residuals (bottom) ($\bar{v}$ = 0.777 cm$^3$ g$^{-1}$). Left: continuous c(s) distribution from sedimentation-velocity data at 60k rpm returning s = 1.392 S, s20,w = 1.728 S, f/f0 = 1.247 and mw = 15,066 Da (4.6 × monomer mass) at 95% confidence level. Conditions: 150 µM peptide concentration, PBS (pH 7.4). Right: sedimentation-equilibrium data (top, dots) and fitted single-ideal species model curves at 30k (blue), 33k (red) and 36k (purple) rpm. The fit returns a mass of 14,430 Da (4.4 × monomer mass, 95% confidence limits 14328 – 14,430). Bottom: residuals for the above fits using the same colour scheme as above. Conditions: 70 µM peptide concentration, PBS (pH 7.4).*

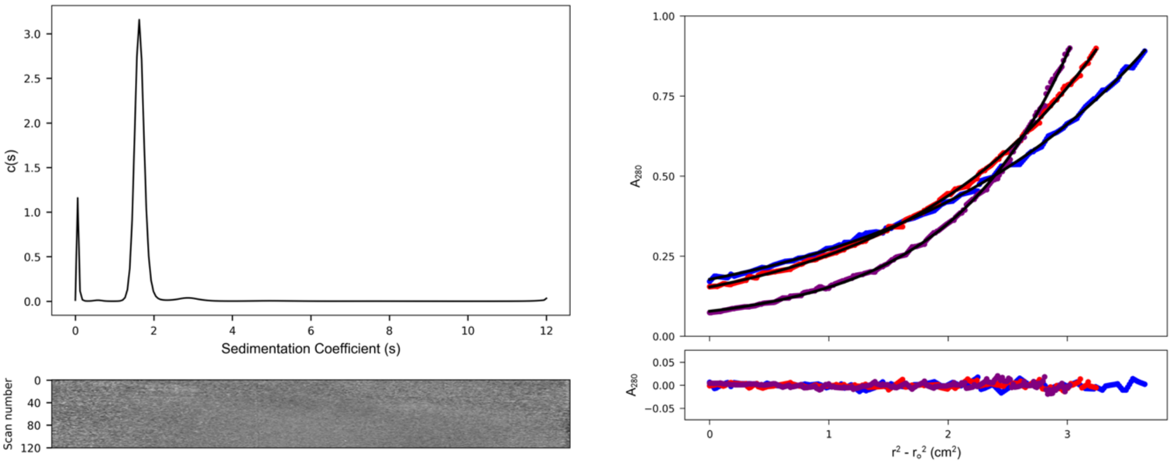


*Figure S6.8 CC-Hex-II AUC data and fits (top) and residuals (bottom) ($\bar{v}$ = 0.777 cm$^3$ g$^{-1}$). Left: continuous c(s) distribution from sedimentation-velocity data at 60k rpm returning s = 1.639 S, s20,w = 1.708 S, f/f0 = 1.245 and mw = 19,677 Da (6.0 × monomer mass) at 95% confidence level. Conditions: 150 µM peptide concentration, PBS (pH 7.4). Right: sedimentation-equilibrium data (top, dots) and fitted single-ideal species model curves at 23k (blue), 26k (red) and 33k (purple) rpm. The fit returns a mass of 18,270 Da (5.6 × monomer mass, 95% confidence limits 18,202 – 18,344). Bottom: residuals for the above fits using the same colour scheme as above. Conditions: 70 µM peptide concentration, PBS (pH 7.4).*

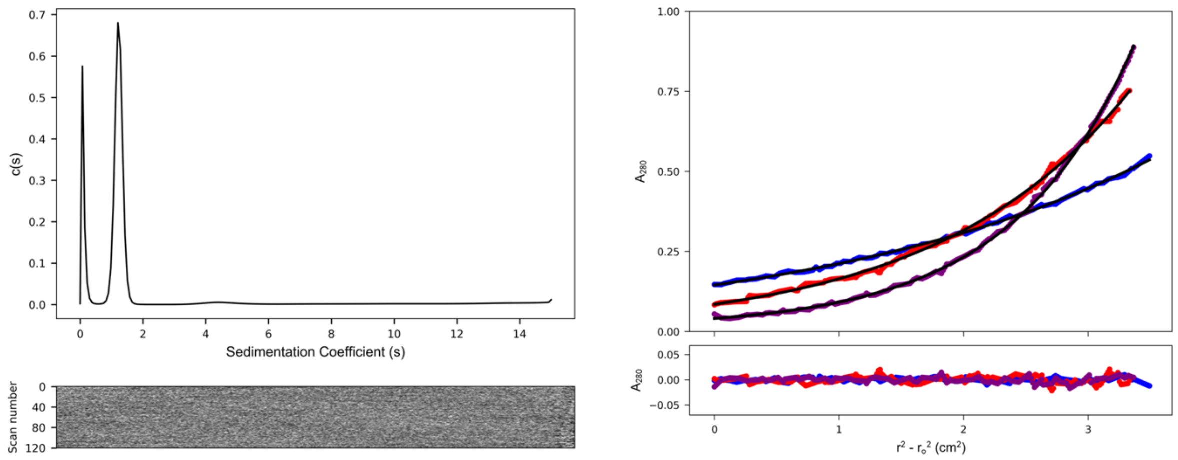


*Figure S6.9 CC-Hex-LL-KgEb AUC data and fits (top) and residuals (bottom) ($\bar{v}$ = 0.777 cm$^3$ g$^{-1}$). Left: continuous c(s) distribution from sedimentation-velocity data at 60k rpm returning s = 1.229 S, s20,w = 1.280 S, f/f0 = 1.233 and mw = 12,590 Da (3.9 × monomer mass) at 95% confidence level. Conditions: 35 µM peptide concentration, PBS (pH 7.4). Right: sedimentation-equilibrium data (top, dots) and fitted single-ideal species model curves at 24k (blue), 32k (red) and 40k (purple) rpm. The fit returns a mass of 12,860 Da (4.0 × monomer mass, 95% confidence limits 12,813 – 12,907). Bottom: residuals for the above fits using the same colour scheme as above. Conditions: 35 µM peptide concentration, PBS (pH 7.4).*

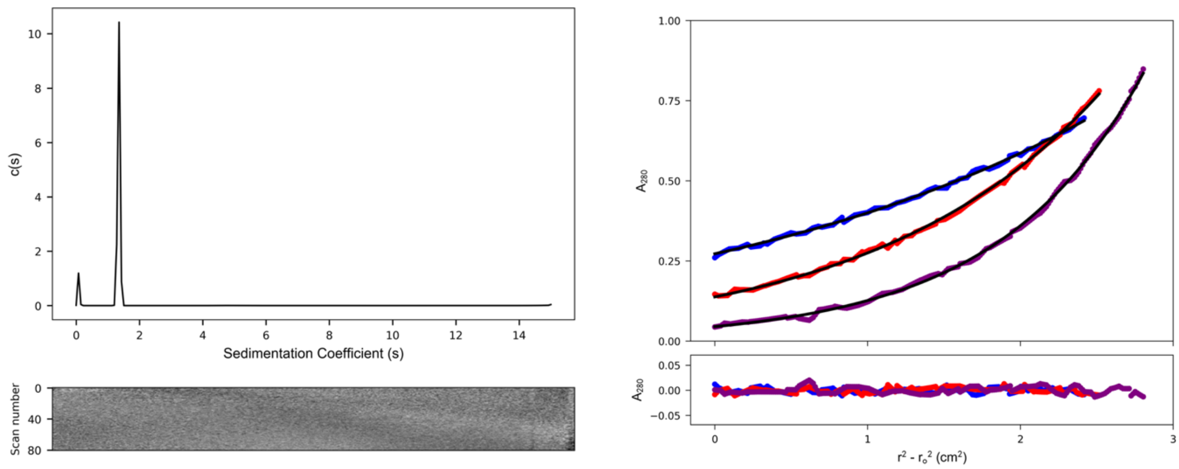


*Figure S6.10 apCC-Tet AUC data and fits (top) and residuals (bottom) ($\bar{v}$ = 0.777 cm$^3$ g$^{-1}$). Left: continuous c(s) distribution from sedimentation-velocity data at 60k rpm returning s = 1.352 S, s20,w = 1.408 S, f/f0 = 1.187 and mw = 13,059 Da (4.0 × monomer mass) at 95% confidence level. Conditions: 150 µM peptide concentration, PBS (pH 7.4). Right: sedimentation-equilibrium data (top, dots) and fitted single-ideal species model curves at 24k (blue), 32k (red) and 40k (purple) rpm. The fit returns a mass of 13,360 Da (4.1 × monomer mass, 95% confidence limits 13,303 – 13,414). Bottom: residuals for the above fits using the same colour scheme as above. Conditions: 70 µM peptide concentration, PBS (pH 7.4).*

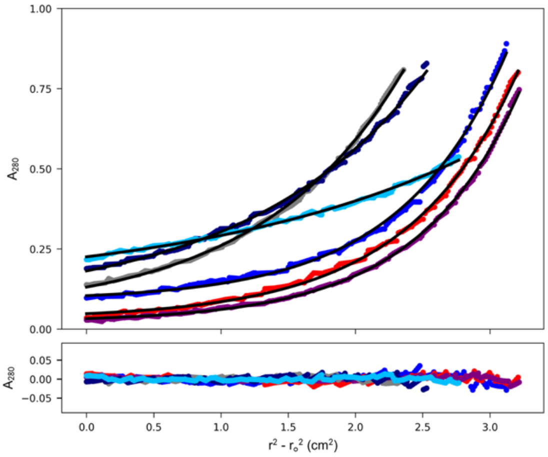


*Figure S6.11 CC-Hex-L24E AUC data and fits (top) and residuals (bottom) ($\bar{\upsilon}$ = 0.768 cm$^3$ g$^{-1}$) at various pH. Sedimentation-equilibrium data (top, dots) and fitted single-ideal species model curves at 36k rpm. Representative scans are shown for pH 3 (dark blue), pH 4 (red), pH 5 (purple), pH 6 (gray), pH 7 (navy blue) and pH 7.4 (light blue). Bottom: residuals for the above fits using the same colour scheme as above. Conditions: 70 µM peptide concentration, 137 mM NaCl, 2.7 mM KCl and a buffer mixture comprising varying ratios of citric acid and $Na_2HPO_4$ to give appropriate pH, with combined buffer concentrations totalling 50-100 mM.*

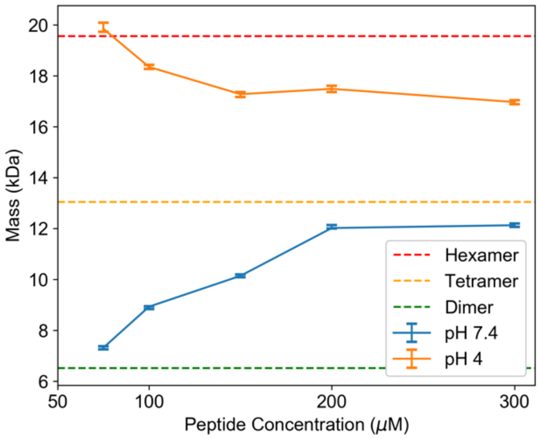


*Figure S6.12 CC-Hex-L24E AUC ($\bar{\upsilon}$ = 0.768 cm$^3$ g$^{-1}$) at various concentrations. Sedimentation-equilibrium data collected from absorbance scans obtained at 4krpm intervals from 20– 48k rpm fitted to single-ideal species speeds. Conditions are as follows: at pH 7.4, 137 mM NaCl, 2.7 mM KCl, 8.2 mM $Na_2HP0_4$ and 1.8 mM $KH_2P0_4$; at pH 4, 30.73 mM citric acid/38.55 mM Na2HPO4 buffer, 137 mM NaCl and 2.7 mM KCl. Error bars represent 95% confidence limits obtained from Monte Carlo.*

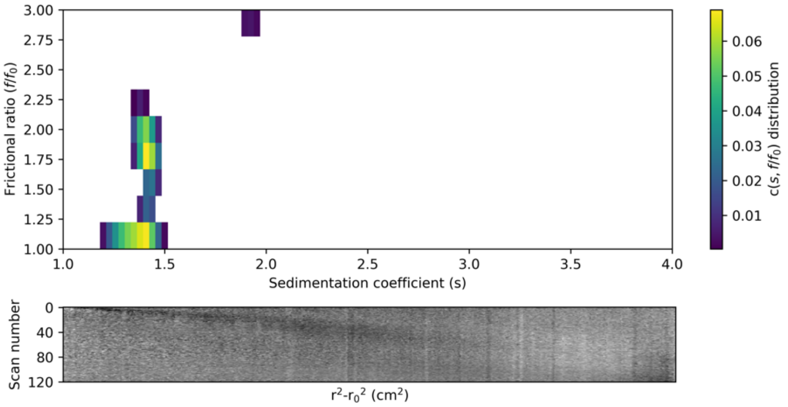


*Figure S6.13 CC-Hex-L24E AUC data and fits (top) and residuals (bottom) ($\overline{\nu}$ = 0.768 cm$^3$ g$^{-1}$). Continuous c(s,f/$f_0$) distribution from sedimentation-velocity data at 60k rpm returning the following integrated peaks: peak 1 and 2 represent 72% of the total loading concentration with peak 1 giving an average sw = 1.372 S, average f/f0 = 1.022 and mw = 11,192 Da (3.4 × monomer mass) at 95% confidence level; peak 2 has an average sw = 1.393 S, average f/f0 = 1.722 and mw = 25,065 Da (7.7 × monomer mass) at 95% confidence level; peak 3 represent 4% of the total loading concentration and sw = 1.908 S, f/f0 = 2.950 and mw = 90,054 Da (27.61 × monomer mass) at 95% confidence level. Conditions: 150 µM peptide concentration, 30.73 mM citric acid/38.55 mM Na2HPO4 buffer, 137 mM NaCl and 2.7 mM KCl (pH 4).*

### Section 7 Crystallography Tables

*Table S2 Crystallisation conditions used to obtain the structures discussed in this article. *These are the final concentrations based on a 1:1 dilution with the peptide solution.*

| Sequence | Crystallisation conditions* | Molecular dimensions screen |
|---|---|---|
| CC-Hex-L24D | 50 mM Tris buffer and 1 M ammonium sulphate at pH 8.5 | Structure screen 1 and 2 C8 |
| CC-Hex-L24E (tetramer) | 50 mM Tris buffer and 1 M ammonium sulphate at pH 8.5 | Structure screen 1 and 2 C8 |
| CC-Hex-L24E (hexamer) | 100 mM NaCl, 50 mM MES buffer and 10% w/v PEG 2000 MME at pH 6.0 | Proplex A9 |
| CC-Hex-L24K | 50 mM Tris buffer and 750 mM ammonium sulphate at pH 8.0 | Proplex G1 |
| CC-Hex-L24Dab | 800 mM $Na_2HPO_4/K_2HPO_4$ at pH 6.5 | Proplex G10 |
| CC-Hex-L24Nle | 50 mM MES buffer, 800 mM ammonium sulphate and 5% v/v 1,4-Dioxane at pH 6.5 | Structure screen 1 and 2 F11 |
| CC-Hex-L24H | 50 mM MES buffer, 25 mM cesium chloride and 15% v/v Jeffamine M-600 at pH 6.5 | Structure screen 1 and 2 F12 |
| CC-Hex-LL | 50 mM Tris buffer and 600 mM $Na_2/K_2$ tartrate at pH 8.0 | Proplex H5 |
| CC-Hex-II | 50 mM SPG buffer (Succinic Acid, sodium phosphate monobasic monohydrate, Glycine) and 12.5 % w/v PEG 1500 at pH 8.0 | Pact A5 |
| CC-Hex-KgEb_var | 100 mM potassium sodium tartrate tetrahydrate and 10% w/v PEG 3350 pH 7.4 | PEG/Ion HT D1 (Hampton Research) |
| CC-Hex-LL-KgEb | 630 mM ammonium sulphate, 100 mM lithium sulphate, 50 mM Tris buffer at pH 8.5 | JCSG E4 |
| apCC-Tet | 50 mM HEPES and 2.15 M NaCl at pH 7.5 | Structure screen 1 and 2 F6 |

*Table S3 X-ray crystallography data collection and processing, and model refinement.*

| | CC-Hex-L24D | CC-Hex-L24E (tetramer) | CC-Hex-L24E (hexamer) |
|---|---|---|---|
| PDB ID code | 6Q5H | 6Q5I | 6Q5J |
| Wavelength (Å) | 0.97628 | 0.97628 | 0.97628 |
| Resolution range (Å) | 15.87-1.2 [15.87-5.37] (1.23-1.20) | 17.38-1.76 [17.38-7.87] (1.81-1.76) | 46.32-1.69 [46.32-7.56] (1.75-1.69) |
| Space group | P 61 2 2 | P 61 2 2 | P 42 21 2 |
| Unit cell lenghts (Å) | 70.20 70.20 35.58 | 69.07 69.07 35.49 | 55.07 55.07 138.94 |
| Unit cell angles (°) | 90.0 90.0 120.0 | 90.0 90.0 120.0 | 90.0 90.0 90.0 |
| Total reflections | 603506 [6907] (41713) | 192879 [2045] (14958) | 610321 [7200] (59742) |
| Unique reflections | 16641 [228] (1215) | 5281 [75] (382) | 24910 [365] (2406) |
| Multiplicity | 36.3 [30.3] (34.3) | 36.5 [27.3] (39.2) | 24.5 [19.7] (24.8) |
| Completeness (%) | 99.7 [96.3] (99.9) | 99.5 [91.2] (100.0) | 100 [100] (100) |
| Mean I/sigma(I) | 17.2 [73.2] (1.1) | 12.9 [52.4] (1.0) | 10.6 [35.1] (2.6) |
| Wilson B-factor (Å$^2$) | 16.17 | 29 | 22 |
| R-merge(I) | 0.098 [0.039] (4.049) | 0.205 [0.051] (4.738) | 0.193 [0.040] (1.004) |
| R-meas(I) | 0.099 [0.039] (4.110) | 0.208 [0.052] (4.800) | 0.197 [0.042] (1.025) |
| CC1/2 | 1 [1] (0.728) | 0.999 [0.999] (0.661) | 1 [1] (0.546) |
| Reflections used in refinement | 16617 (1615) | 5264 (504) | 24854 (2376) |
| Reflections used for R-free | 842 (81) | 259 (25) | 1205 (108) |
| R-work | 0.151 (0.375) | 0.1730 (0.3499) | 0.2161 (0.408) |
| R-free | 0.180 (0.378) | 0.2060 (0.3534) | 0.2579 (0.361) |
| Number of non-hydrogen atoms | 504 | 447 | 1594 |
| macromolecules | 430 | 400 | 1341 |
| ligands | 18 | 18 | 43 |
| Protein residues | 54 | 58 | 177 |
| RMS(bonds) | 0.019 | 0.0166 | 0.014 |
| RMS(angles) | 2.06 | 1.8906 | 1.76 |
| Ramachandran favored (%) | 98 | 96 | 100 |
| Ramachandran allowed (%) | 1.9 | 4.2 | 0 |
| Ramachandran outliers (%) | 0 | 0 | 0 |
| Rotamer outliers (%) | 7.5 | 0 | 0.83 |
| Clashscore | 8.65 | 7 | 4.17 |
| Average B-factor | 44.71 | 39.25 | 28.72 |
| macromolecules | 45.19 | 37.48 | 26.54 |
| ligands | 38.65 | 64.41 | 45.43 |
| solvent | 43.04 | 48.05 | 39.16 |
| Number of TLS groups | | 2 | 6 |

| | CC-Hex-L24K | CC-Hex-L24Dab | CC-Hex-L24Nle |
|---|---|---|---|
| PDB ID code | 6Q5K | 6Q5M | 6Q5N |
| Wavelength (Å) | 0.9200 | 0.97833 | 0.97935 |
| Resolution range (Å) | 24.77-1.65 [24.77-9.04] (1.68-1.65) | 35.69-1.50 [35.69-8.22] (1.53-1.50) | 47.92-2.0 [47.92-8.94] (2.05-2.00) |
| Space group | P 61 2 2 | P 61 2 2 | P 21 21 21 |
| Unit cell lenghts (Å) | 69.28 69.28 35.44 | 71.38 71.38 35.40 | 54.19 59.29 143.77 |
| Unit cell angles (°) | 90 90 120 | 90 90 120 | 90 90 90 |
| Total reflections | 107785 [767] (5050) | 38423 [256] (1909) | 291324 [2899] (21713) |
| Unique reflections | 6135 [53] (269) | 8775 [256] (433) | 32110 [375] (2340) |
| Multiplicity | 17.6 [14.5] (18.8) | 4.4 [3.9] (4.4) | 9.1 [7.7] (9.3) |
| Completeness (%) | 96.5 [95.4] (90.2) | 98.9 [89.3] (100) | 99.8 [85.8] (99.9) |
| Mean I/sigma(I) | 20.4 [50.4] (2.7) | 12.0 [43.1] (1.1) | 13.0 [27.2] (1.5) |
| Wilson B-factor (Å$^2$) | 25.21 | 21.63 | 36.52 |
| R-merge(I) | 0.0701 [0.029] (1.356) | 0.053 [0.015] (1.173) | 0.105 [0.046] (1.730) |
| R-meas(I) | 0.073 [0.030] (1.393) | 0.060 [0.018] (1.331) | 0.111 [0.049] (1.833) |
| CC1/2 | 0.999 [1.000] (0.682) | 1.0 [1.0] (0.703) | 0.999 [1.000] (0.511) |
| Reflections used in refinement | 6134 (556) | 8322 (863) | 32020 (3151) |
| Reflections used for R-free | 313 (27) | 447 (27) | 1558 (153) |
| R-work | 0.184 (0.3008) | 0.202 (0.3327) | 0.234 (0.340) |
| R-free | 0.226 (0.3007) | 0.233 (0.3339) | 0.274 (0.353) |
| Number of non-hydrogen atoms | 472 | 456 | 2917 |
| macromolecules | 421 | 399 | 2698 |
| ligands | 19 | 18 | 108 |
| Protein residues | 53 | 52 | 368 |
| RMS(bonds) | 0.018 | 0.016 | 0.05 |
| RMS(angles) | 2.01 | 1.84 | 1.76 |
| Ramachandran favored (%) | 98 | 100 | 99 |
| Ramachandran allowed (%) | 2 | 0 | 0.3 |
| Ramachandran outliers (%) | 0 | 0 | 0.6 |
| Rotamer outliers (%) | 2.6 | 0 | 1.7 |
| Clashscore | 10.81 | 3.48 | 7.1 |
| Average B-factor | 34.27 | 32.04 | 45.3 |
| macromolecules | 32.88 | 29.75 | 44.32 |
| ligands | 45.15 | 54.45 | 60.26 |
| solvent | 46.12 | 45.14 | 54.59 |
| Number of TLS groups | 2 | | |

| | CC-Hex-L24H | CC-Hex-LL | CC-Hex-II |
|---|---|---|---|
| PDB ID code | 6Q5L | 6Q5O | 6Q5P |
| Wavelength (Å) | 0.9795 | 0.77999 | 0.9163 |
| Resolution range (Å) | 27.96-1.41 [27.96-6.31] (1.45-1.41) | 27.56-1.0 [27.56-2.71] (1.036-1.0) | 82.61-1.44 [82.61-6.44] (1.491-1.44) |
| Space group | P 21 21 2 | I 2 2 2 | P 63 |
| Unit cell lenghts (Å) | 27.98 30.05 53.87 | 28.87 29.38 55.12 | 62.76 62.76 82.62 |
| Unit cell angles (°) | 90 90 90 | 90 90 90 | 90 90 120 |
| Total reflections | 85130 [1244] (1565) | 163349 [8207] (16449) | 664157 [7736] (63526) |
| Unique reflections | 8784 [136] (452) | 13041 [721] (1287) | 33440 [398] (3324) |
| Multiplicity | 9.7 [9.1] (3.5) | 12.5 [11.4] (12.8) | 19.9 [19.4] (19.1) |
| Completeness (%) | 95.2 [98.7] (70.9) | 100 [100] (100) | 100 [100] (100) |
| Mean I/sigma(I) | 11.0 23.0 (1.0) | 15.5 [61.8] (0.5) | 11.5 [49.1] (1.3) |
| Wilson B-factor (Å$^2$) | 20.19 | 15.78 | 18.22 |
| R-merge(I) | 0.088 [0.104] (0.857) | 0.050 [0.035] (3.861) | 0.146 [0.047] (3.021) |
| R-meas(I) | 0.092 [0.110] (1.008) | 0.052 [0.037] (4.02) | 0.150 [0.049] (3.104) |
| CC1/2 | 0.998 [0.997] (0.492) | 0.999 [0.999] (0.491) | 0.999 [0.999] (0.639) |
| Reflections used in refinement | 8766 (631) | 13041 (1276) | 33408 (3315) |
| Reflections used for R-free | 411 (27) | 686 (87) | 1678 (140) |
| R-work | 0.211 (0.356) | 0.168 (0.412) | 0.165 (0.298) |
| R-free | 0.247 (0.354) | 0.212 (0.452) | 0.171 (0.290) |
| Number of non-hydrogen atoms | 432 | 341 | 1680 |
| macromolecules | 386 | 314 | 1463 |
| ligands | 18 | 1 | 32 |
| Protein residues | 52 | 31 | 188 |
| RMS(bonds) | 0.016 | 0.016 | 0.015 |
| RMS(angles) | 1.85 | 1.78 | 1.58 |
| Ramachandran favored (%) | 100 | 100 | 100 |
| Ramachandran allowed (%) | 0 | 0 | 0 |
| Ramachandran outliers (%) | 0 | 0 | 0 |
| Rotamer outliers (%) | 0 | 6.5 | 1.5 |
| Clashscore | 5.97 | 10.13 | 6.71 |
| Average B-factor | 32.61 | 20.59 | 25.81 |
| macromolecules | 29.55 | 19.07 | 24.06 |
| ligands | 81.48 | 42.25 | 44.19 |
| solvent | 43.43 | 38.15 | 36.51 |
| Number of TLS groups | 2 | | 6 |

| | CC-Hex-KgEb_var | CC-Hex-LL-KgEb | apCC-Tet |
|---|---|---|---|
| PDB ID code | 6Q5Q | 6Q5R | 6Q5S |
| Wavelength (Å) | 0.9795 | 0.9686 | 0.97628 |
| Resolution range (Å) | 20.47-1.08 [20.47-3.42] (1.14-1.08) | 42.34-1.61 [42.34-7.20] (1.65-1.61) | 48.96-1.89 [48.96-8.45] (1.94-1.89) |
| Space group | I 2 2 2 | P 1 | P 41 21 2 |
| Unit cell lenghts (Å) | 28.57 29.35 55.72 | 45.63 46.75 48.16 | 58.49 58.49 89.47 |
| Unit cell angles (°) | 90 90 90 | 89 62 82 | 90 90 90 |
| Total reflections | 132600 [4230] (17765) | 145061 [1718] (10552) | 328587 [3639] (24695) |
| Unique reflections | 10348 [353] (1461) | 43409 [500] (3130) | 13052 [196] (940) |
| Multiplicity | 12.8 [12.0] (12.2) | 3.3 [3.4] (3.4) | 25.2 [18.6] (26.3) |
| Completeness (%) | 99.7 [94.7] (98.8) | 97.1 [99.2] (95.4) | 99.9 [99.2] (99.9) |
| Mean I/sigma(I) | 17.5 [56.2] (3.2) | 4.8 [8.8] (2.6) | 15.1 [45.7] (1.1) |
| Wilson B-factor (Å$^2$) | 9.42 | 14.43 | 35.65 |
| R-merge(I) | 0.074 [0.040] (0.706) | 0.191 [0.239] (1.095) | 0.146 [0.044] (3.527) |
| R-meas(I) | 0.077 [0.042] (0.737) | 0.228 [0.286] (1.308) | 0.149 [0.045] (3.596) |
| CC1/2 | 0.999 [0.999] (0.943) | 0.945 [0.927] (0.503) | 0.999 [1.000] (0.553) |
| Reflections used in refinement | 10315 (997) | 43407 (4130) | 13004 (1263) |
| Reflections used for R-free | 495 (40) | 2076 (192) | 616 (53) |
| R-work | 0.138 (0.2720) | 0.196 (0.218) | 0.198 (0.291) |
| R-free | 0.149 (0.2416) | 0.237 (0.273) | 0.232 (0.324) |
| Number of non-hydrogen atoms | 288 | 3135 | 995 |
| macromolecules | 240 | 2832 | 909 |
| ligands | 14 | 27 | 10 |
| Protein residues | 34 | 367 | 124 |
| RMS(bonds) | 0.019 | 0.006 | 0.007 |
| RMS(angles) | 1.81 | 1 | 1 |
| Ramachandran favored (%) | 100 | 100 | 100 |
| Ramachandran allowed (%) | 0 | 0 | 0 |
| Ramachandran outliers (%) | 0 | 0 | 0 |
| Rotamer outliers (%) | 0 | 0.78 | 2.5 |
| Clashscore | 3.82 | 8.87 | 6.73 |
| Average B-factor | 16.37 | 23.1 | 41.11 |
| macromolecules | 13.91 | 21.54 | 40.26 |
| ligands | 22.02 | 47.79 | 67.38 |
| solvent | 31.43 | 36.63 | 47.83 |
| Number of TLS groups | | 12 | 1 |
